# Machine learning assisted classification of cell and brain penetrating peptides

**DOI:** 10.1101/2025.08.26.672010

**Authors:** Vito Kontrimas, Gabriele C. Sosso, Robert Dallmann, Sebastien Perrier

## Abstract

Crossing the blood–brain barrier (BBB) remains a major obstacle for central nervous system therapeutics. Short peptides have emerged as promising vectors, including cell-penetrating peptides (CPPs) and brain-penetrating peptides (BPPs). However, the structural and physicochemical features that distinguish CPPs from BPPs remain poorly understood, limiting rational design. Here, we compiled a curated dataset of 490 peptides, encompassing CPPs, BPPs, and non-BPP controls, and systematically analysed their amino acid composition, sequence distribution, and physicochemical descriptors. BPPs were found to exhibit a more balanced distribution of cationic, polar, and hydrophobic residues compared to CPPs, which were enriched in contiguous arginine and lysine blocks. Physicochemical analysis revealed that BPPs had lower charge density, greater stability, and reduced aromaticity relative to CPPs. Dimensionality reduction confirmed BPPs occupy an intermediate chemical space between CPPs and non-BPPs. Machine learning classification, particularly with Extra Trees models, achieved strong performance in discriminating peptide classes, with charge, instability index, and aromaticity identified as the most predictive features. These findings suggest that BBB penetration is not a simple extension of cell penetration but requires finely tuned physicochemical properties. This study provides mechanistic insights into BPP design and highlights machine learning as a valuable tool for engineering next-generation BBB-penetrating peptides and peptide-mimetic materials.

## Introduction

Traversing the blood-brain barrier (BBB) remains a significant challenge, limiting therapeutic strategies available ^1^. Small peptides have been explored to address this challenge, primarily via the trojan-horse approach to deliver hydrophilic, polar compounds, larger proteins or even genetic materials across the cellular barriers, such as the BBB^2,3^. These peptides typically exhibit cationic and hydrophobic properties, characteristic of amphiphilic peptides^4,5^, and are mostly linear, suggesting they have simple conformational structures such as alpha helices or beta sheets ^2,3^. However, the specific properties that enable certain peptides to successfully cross the blood-brain barrier are still under debate. Not all cell-penetrating peptides (CPPs)^4^ can cross the blood-brain barrier ^5,6^, yet precise compositional and structural differences between CPPs and BPPs still remain unclear. This knowledge gap jeopardises the rational development of novel peptides or synthetic peptide-like compounds, that would show blood-brain barrier permeability properties ^7^. Over the past decade, the use of computational approaches to study biological proteins and peptides has steadily increased ^8–10^, with advancements in machine-learning research, data mining and access to computing power transforming how researchers analyse biological data ^11,12^. For example, such methods enable the extraction of meaningful data patterns with potential translational applicability. Arif et al. showed that the Gradient Boost Decision Tree (GBDT) algorithm using simple descriptors, such as amino acid composition, peptide length, and other physicochemical properties of amino acids, was able to successfully predict cell-penetrating peptides ^13^. Another study by Hsueh et al.^14^ also used an ensemble of decision tree algorithms to classify CPPs. The important features found in CPPs were then used to engineer a novel peptide that targeted retinal pigment epithelium cell line (ARPE-19) and outperformed the transactivator of transcription (TAT) CPP. Gu et al. showed the classification of brain-penetrating peptides using various decision tree algorithms with an excellent performance ^7^. A more recent meta-analysis study comparing CPPs and BPPs proposed that most CPPs don’t cross the blood-brain barrier, suggesting distinct physicochemical properties between these two groups ^15^.

These data-driven research methods offer better insights into peptide structures and can assist in novel compound synthesis. In this study, we hypothesised that identifiable physicochemical differences exist between CPPs, BPPs and peptides that cannot cross the BBB which could be analysed through machine learning. First, we collected and refined the datasets from literature and public datasets comprising CPPs, BPPs and peptides that cannot cross the BBB (non-BPPs). The well-known physicochemical property descriptors such as amino acid composition, distribution and physicochemical properties of amino acids were extracted for each class of the peptides and data analysis was explored. Later, we employed decision tree algorithms to classify these peptides and measure the algorithm’s performance. Finally, we extracted the important features that distinguished these peptides from one another.

## Methodology

### Peptide database

The database of brain-penetrating peptides (BPPs), cell-penetrating peptides (CPPs), and non-brain-penetrating peptides (non-BPPs) was collected from public datasets CPPSite2.0, BrainPeps, and B3Pdb and literature. Only the peptides with natural amino acids were selected, excluding any peptides containing chemical modifications. The dataset comprised a mixture of experimental conditions (both in vitro and in vivo) and natural and synthetic peptides. The summary of the dataset peptide frequencies is displayed in Table 1.

**Table 1.** List of peptide groups and their peptide frequencies in the dataset.

|  | Number of peptides |
| --- | --- |
| BPPs | 191 |
| CPPs | 273 |
| Non-BPPs | 26 |

### Amino acid analysis

The amino acid frequency in BPPs, CPPs and non-BPPs was calculated using Python and Bio.SeqUtils.ProtParam module (Biopython). Briefly, the total count of each amino acid was calculated in an individual category of peptides and then divided by the total number of amino acids in a group, with the representative formula below:

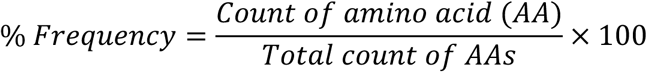

Statistical analysis was performed using pairwise Welch’s t-test with Bonferroni correction. Welch’s t-test was selected due to an imbalanced dataset, as described previously.

### Physicochemical feature extraction

The key physicochemical descriptors of the peptides were extracted using modlAMP (version 4.2.3). Specifically, we calculated the molecular weight (Mw), charge density, isoelectric point (pI), hydrophobic ratio, Boman index, aromaticity, aliphatic index, instability index, total charge, and length of the peptides using the modlamp.descriptors.GlobalDescriptor function. The modlAMP descriptors are summarised in Table 2.

**Table 2.** Summary of modlAMP descriptors. Adapted from modlAMP by Müller et al.^16^.

| Full descriptor name | Descriptor function | Description |
| --- | --- | --- |
| Peptide Sequence Length | `length()` | Total amino acid count in the peptide sequence |
| Molecular Weight | `calculate_MW` | Molecular weight of the peptide |
| Charge | `calculate_charge()` | Calculates the overall charge of the peptide sequence at specified pH. |
| Charge density | `charge_density()` | Calculates charge per molecular weight |
| Isoelectric point | `isoelectric_point()` | Calculates isoelectric point of each peptide sequence based on pKa values of amino acids |
| Instability index | `instability_index()` | Calculates instability index |
| Aromaticity | `aromaticity()` | Computes relative frequencies of tyrosine (Tyr), tryptophan (Trp) and phenylalanine (Phe). |
| Aliphatic index | `aliphatic_index()` | Calculates aliphatic index |
| Boman index | `boman_index()` | Computes boman index |
| Hydrophobic ratio | `hydrophobic_ratio()` | Relative frequencies of A,C,F,I,L,M & V amino acids. |

### Cationic charge distribution analysis

The peptide amino acid sequences were split into N-terminus, C-terminus and core of the peptide. Then, the cationic charge, charge density and GRAVY score were calculated using modlAMP descriptors as described before.

### Peptide amino acid sequence logo maps

The probability distributions of amino acids at each position of the peptide sequences were calculated and visualised using WebLogo 3.0 sequence logo generator.

### Dimensionality reduction

The dimensionality reduction was calculated for CPPs, BPPs and non-BPPs using Python to assess data clustering. Briefly, the principal component analysis (PCA) from the scikit ^17^ library, t-distributed Stochastic Neighbour Embedding (t-SNE) from the scikit library, and Uniform Manifold Approximation and Projection (UMAP) from the umap-learn ^18^ library were used. Before analysis, the data was standardised using StandardScaler from the scikit learn library. The PCA was used to reduce the data to two principal components, and the explained variance ratio was calculated for each dimension. Likewise, the UMAP was used to reduce the data to two dimensions using default variables (n_components=2). The t-SNE was performed with n_components=2 variable and perplexity set to 30, 60 or 100. The cluster scores were evaluated using Silhouette scores ^19^ using the scikit library. The statistical analysis was performed on the principal component 1 (PC1) of the PCA. The data was plotted using Seaborn and Matplotlib Python libraries.

### Machine learning analysis

#### Data pre-processing

Before the machine learning classification, the BPP, CPP and non-BPP dataset was pre-processed to contain peptide identifier labels and matching descriptors (modlAMP explained above). The data was then split into descriptors (X) and labels (y) using the pandas library. Consequently, data was subdivided into X_train, y_train, X_test and y_test subsets using the scikitlearn train_test_split function with 70-30 split respectively and stratification to address the imbalanced class distribution. The X_train and y_train were then set as inputs for training the classifiers and X_test and y_test to assess their prediction performance.

#### Classifier selection and optimisation

Random Forest, XGBoost, and Extra Trees classifiers were employed to perform peptide classification. Each model was subjected to a unique pipeline containing a scaler (MinMaxScaler was used to normalise descriptors to [0,1] range) and feature selection, which selected the ten most important features based on ANOVA F-values (calculated using SelectKBest) and the classifier function. Each pipeline was then subjected to the hyperparameter optimisation step using the GridSearchCV function. The grid was optimised for n_estimators [10, 100, 500] and max_features [5, 10, 20, 30] parameters. The performance of the classifiers was then evaluated using Repeated stratified K-fold cross-validation with five splits and repeated 3 times. Then, the performance metrics of the classifiers were evaluated. Specifically, F1 score and accuracy were used.

#### Classifier model validation and performance evaluation

The accuracy and F1 scores of the classifiers were used to identify the best-performing classification model. Subsequently, the model was further validated with the independent test dataset. Then, to identify which descriptors (features) influenced the model the most, permutation feature importances and Shapley Additive explanations (SHAP) values were performed, scoring the features that contributed the most to the model’s output.

The confusion matrix was calculated using true positives (TPs), true negatives (TNs), false positives (FPs), and false negatives (FNs). F1-scores were calculated using the following formulas:

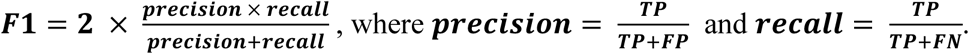

Accuracy scores were calculated as:

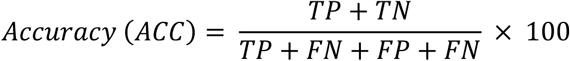

## Results and Discussion

### Amino acid composition analysis of BPPs, CPPs and non-BPPs

The composition of membrane-active peptides such as antimicrobial peptides (AMPs) or CPPs have been documented to contain higher frequencies of cationic or hydrophobic amino acid (AA) residues ^20–22^ which is important for electrostatic interaction with anionic extracellular cell membrane proteoglycans^23,24^. To investigate whether a similar pattern is observed within the peptide database in this study, the amino acid composition and frequency was first analysed (**Figure 1**).

**Figure 1.**
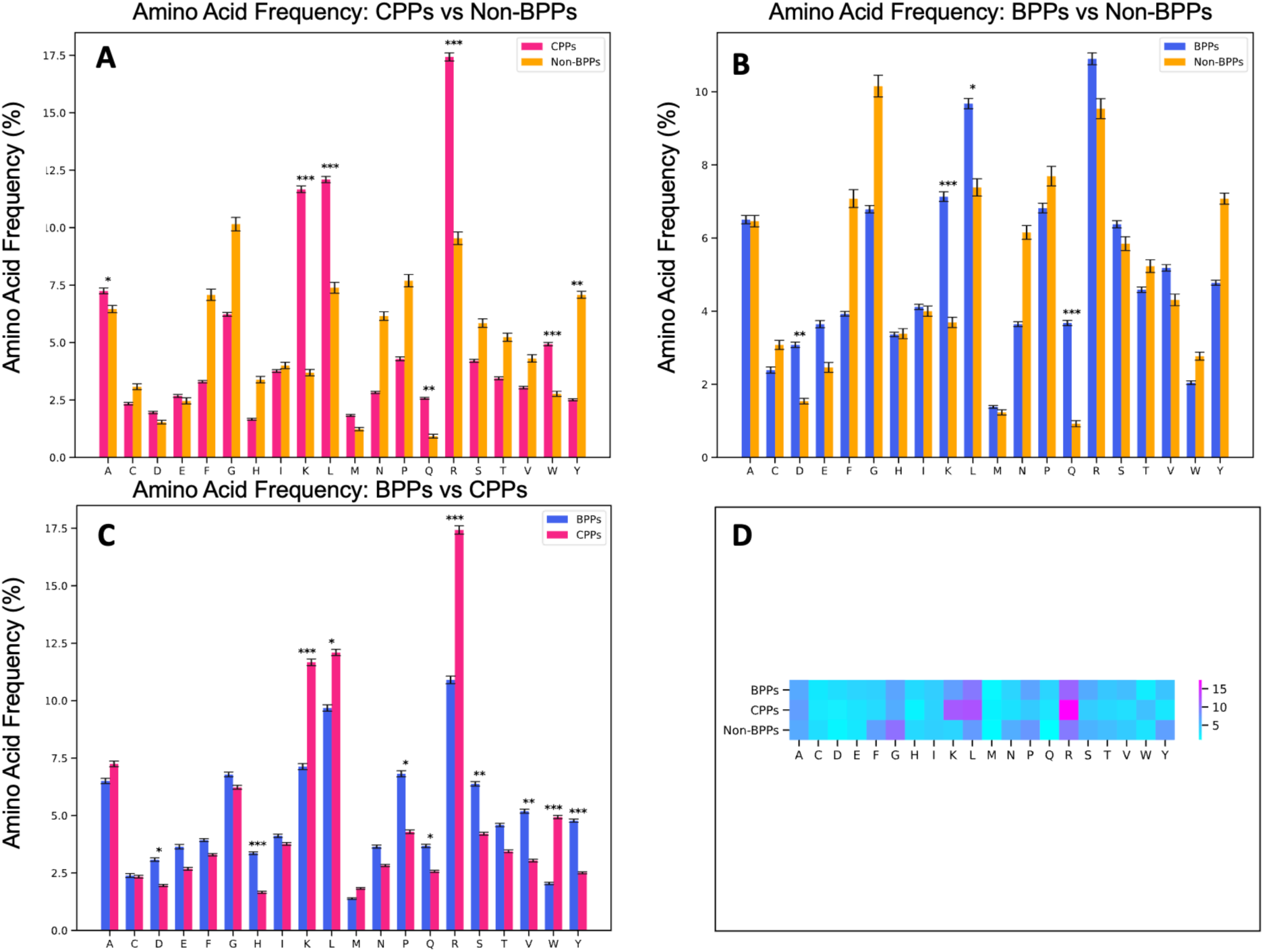
Amino acid distribution in BPPs, CPPs and non-BPPs expressed as percentage frequency comparison and analysis. Comparison of CPPs vs Non-BPPs (A), BPPs vs Non-BPPs (B), BPPs vs CPPs (C) and heatmap summary plot of all peptide classes (D). Data are presented as mean ± standard deviation. Statistical significance was determined using one-way ANOVA followed by Tukey’s post hoc test. *: p < 0.05; **: p < 0.01; ***: p < 0.001.

The results showed that both CPPs and BPPs peptide groups had markedly higher frequencies of cationic amino acids such as lysine (K), hydrophobic leucine (L) and hydrophilic glutamine (Q), in comparison to non-brain-penetrating peptides (non-BPPs) class (**Figure 1A–C**). Additionally, CPPs, unlike BPPs, had higher frequency of hydrophobic amino acids such as alanine (A), tryptophan (W) and tyrosine (Y) as well as cationic arginine (R) within their sequence when compared to non-BPPs. On the other hand, comparing BPPs to non-BPPs indicated a higher content of negatively charged glutamic acid (E) amino acid residues in non-BPPs.

Interestingly, there were also notable differences between CPPs and BPPs. Whilst CPPs were clearly showing more cationic (R, K) and hydrophobic (L, W) amino acids, the BPPs suggested higher content of hydrophobic proline (P) and valine (V), polar serine (S) and histidine (H), amphipathic tyrosine (Y) and anionic aspartic acid (D). Consistent with literature, our results indicated that both BPPs and CPPs indeed both have cationic and amphipathic properties within a limit of our tested dataset. This also correlates with in-vitro studies where amphiphilicity reduces haemolysis and, therefore, increases biocompatibility without compromising the cell penetration properties ^25,26^. Interestingly, our results showed that brain-penetrating peptides, whilst similar to CPPs, had more subtle properties, which may contribute to their success in crossing the blood-brain barrier. It was evident that BPPs had a more balanced amino acid composition, encompassing not only cationic charge and hydrophobic properties but also polar amino acids, indicating higher complexity (**Figure 1A–D**).

### Amino acid distribution and probability analysis of BPPs, CPPs and non-BPPs peptides

The amino acid sequence probability distribution, which calculated the probability of amino acids at each position, revealed distinct positional patterns amongst CPPs, BPPs and non-BPPs (**Figure 2 A-C)** It was observed that BPPs a higher probability of sparse distribution of cationic amino acids, whilst CPPs had a higher probability of continuous cationic blocks within the first 20 amino acid block in their sequence (**Figure 2B**). Such cationic amino acid arrangement appeared less apparent in BPPs (**Figure 2A**) and largely absent in non-BPPs (**Figure 2C**). Lysine amino acid seemed to be enriched within the middle part of the peptides, at approximate positions between 10–25 amino acids in both BPPs and CPPs but not in non-BPPs. On the other hand, non-BPPs had a higher probability of neutral and aliphatic amino acids (**Figure 2C**) but lower probability of cationic amino acids. These results indicated that CPPs, BPPs and non-BPPs had differences not only in their amino acid composition but also their distribution along the peptide sequences.

**Figure 2.**
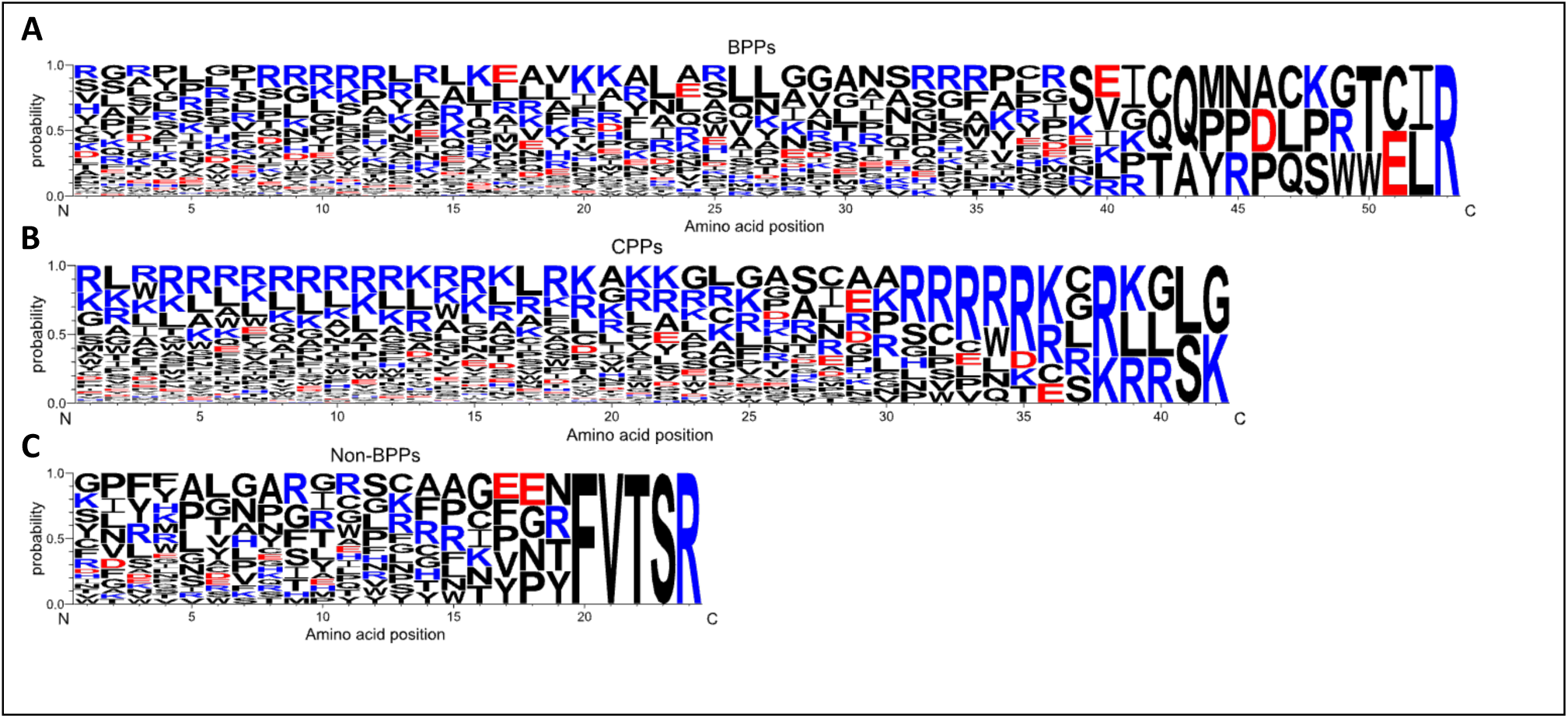
A sequence logo of peptide amino acid distribution in BPPs (A), CPPs (B) and non-BPPs (C), where size is proportional to amino acid probability at the position.

### Theoretical physicochemical properties of BPPs, CPPs and non-BPPs

The physicochemical properties of peptides were analysed using modlAMP peptide descriptors ^16^ and compared amongst BPPs, CPPs and non-BPPs to explore their similarities and differences (**Figure 3**). These results complemented the compositional analysis results outlined in **Figure 1**. The results identified that CPPs and BPPs had similar number of amino acids, as opposed to non-BPPs, which were shorter. There were no observed differences in the relative ratio of hydrophobic amino acids (A, C, F, I, L, M & V) nor aliphatic index between the three peptide groups. Similarly to amino acid composition analysis in **Figure 1**, BPPs were less cationic compared to CPPs, with differences also observed in charge density when compared to CPPs, but not with non-BPPs.

**Figure 3.**
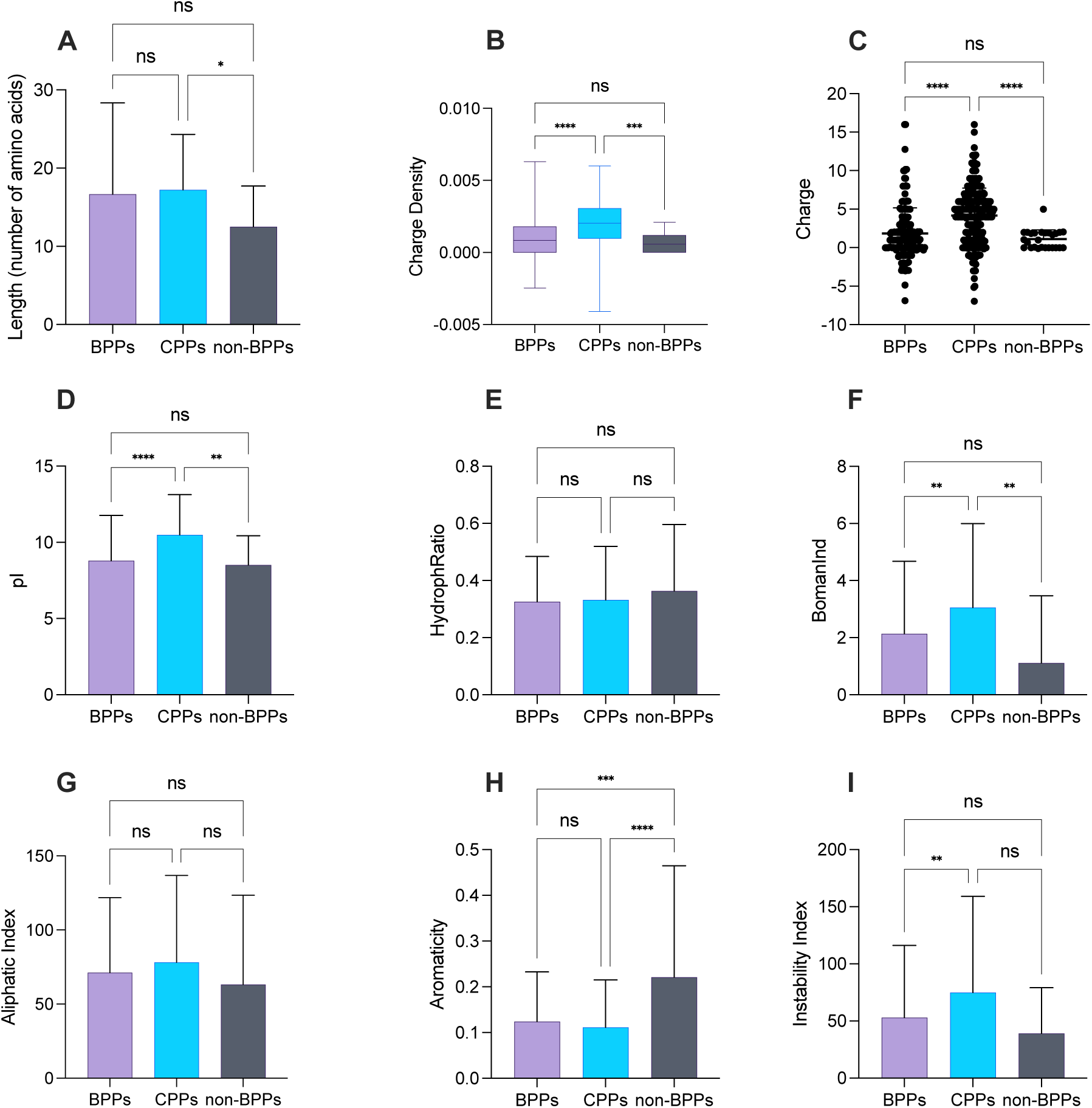
Analysis and comparison of physicochemical properties between brain-penetrating peptides (BPPs), cell-penetrating peptides (CPPs), and non-BPPs. The properties analysed include (A) length, (B) charge density, (C) charge, (D) isoelectric point (pI), (E) hydrophobic ratio, (F) Boman Index, (G) aliphatic index, (H) aromaticity, and (I) instability index.

Isoelectric point (pI) and Boman index (BomanInd) showed a similar pattern of significant differences between the latter groups of peptides. However, both BPPs and CPPs showed significantly lower aromaticity when compared to non-BPPs, suggesting higher probability of steric hindrance in non-BPPs as reported previously ^27,28^. Finally, BPPs had a markedly lower instability index than CPPs, unlike when compared to non-BPPs. The latter suggested that BPPs could exhibit better stability in terms of their bioactivity and enzymatic degradation ^29^.

Several studies have shown that the physicochemical properties of amino acids at the peptide terminals could contribute to membrane pore formations. For example, Strandberg et al.^30^ showed that cationic N-terminus lyses the phospholipid membrane, mimicking vesicles via pore formation, whilst more hydrophobic C-terminus had reduced interaction with membranes. Therefore, the average charge, charge distribution and hydrophobicity were investigated by calculating values for N- and C-terminals as well as the core of the peptide.

As seen in **Figure 4**, splitting the peptides into N-terminus, core and C-terminus showed that BPPs are less cationic than CPPs but not non-BPPs, and more hydrophobic than CPPs at their core. The N-terminus appeared to be less cationic when compared to CPPs and more similar to non-BPPs as observed in **Figure 2** amino acid position probability plots. Comparing BPPs to CPPs, the C-terminus had no difference to CPPs; however, it was more cationic and less hydrophobic than non-BPPs.

**Figure 4.**
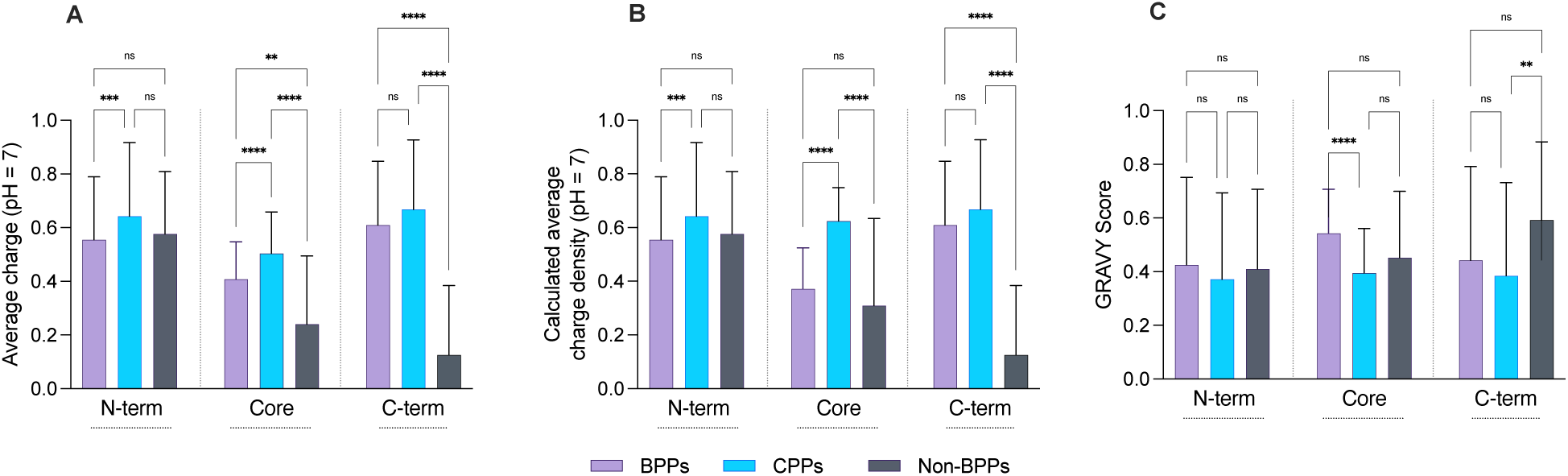
Comparison of peptide charge (A), charge density (B) and Gravy index score (C) at three different positions within a peptide: N-terminus, C-terminus, and the core.

### Dimensionality reduction of descriptors

To investigate whether BPPs, CPPs and non-BPPs could be clustered into distinct groups based on their overall physicochemical properties, we applied several unsupervised dimensionality reduction methods. Unlike manual peptide descriptor comparison, unsupervised machine-learning approach allowed us to investigate whether peptides can be separated into distinct groups when considering their descriptors in their entirety. Here we have used principal component analysis (PCA) ^31,32^, Uniform Manifold Approximation and Projection for Dimension Reduction (UMAP)^18^ and t-distributed Stochastic Neighbour Embedding (t-SNE) ^33^ algorithms. We have used the same descriptor set outlined in the methodology section (Table 2).

The linear dimensionality reduction algorithm (PCA) which identifies largest source of variance in a dataset, showed a lack of separation between BPPs, CPPs and non-BPPs (PC1 = 39.13%, PC2 = 21.51%, **Figure 5A**). Similarly, the non-linear UMAP algorithm showed no distinct clusters, although some local, and sparse clusters were observed when comparing CPPs, BPPs and non-BPPs (**Figure 5B**). A different type of non-linear clustering, t-SNE, with three distinct perplexity parameters, was also unable to separate BPPs, CPPs and non-BPPs (**Figure 5C, 5D, 5E**). To better quantify the dimensionality reduction findings, we calculated a Silhouette clustering (S-clust) ^19^ scores which measured the distances individual points to other points within a group and compared distances to neighbouring groups of points.

**Figure 5.**
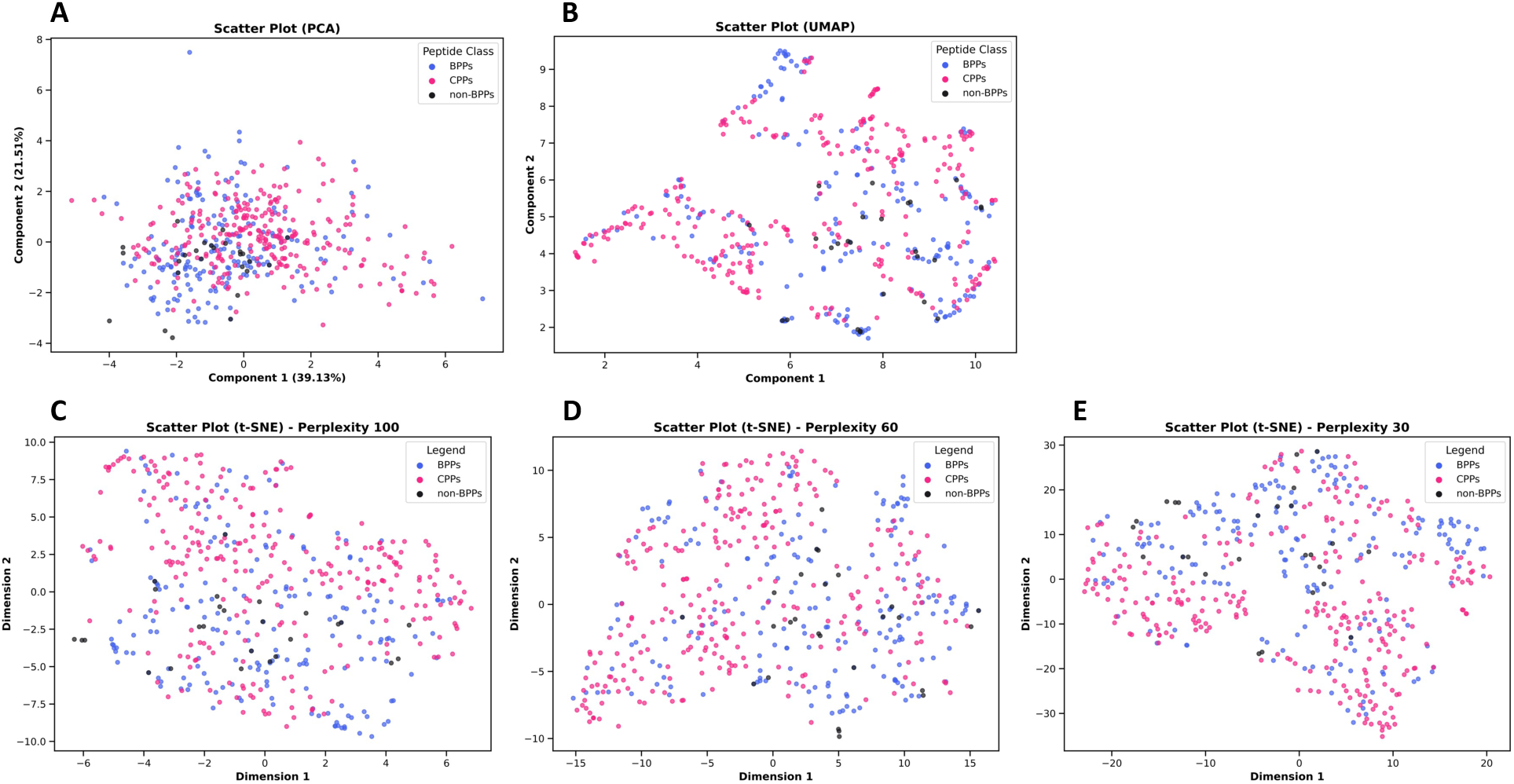
Dimensionality reduction analysis using (A) PCA, (B) t-SNE, and (C) UMAP. Clusters represent CPPs, BPPs, and non-BPPs peptides (**B**) shows Uniform Manifold Approximation and Projection for Dimension Reduction (UMAP) of the same peptide groups. We have also measured t-distributed Stochastic Neighbour Embedding (t-SNE) with variable perplexities of 100 (**C**), 60 (**D**) and 30 (**E**).

**Figure 6.**
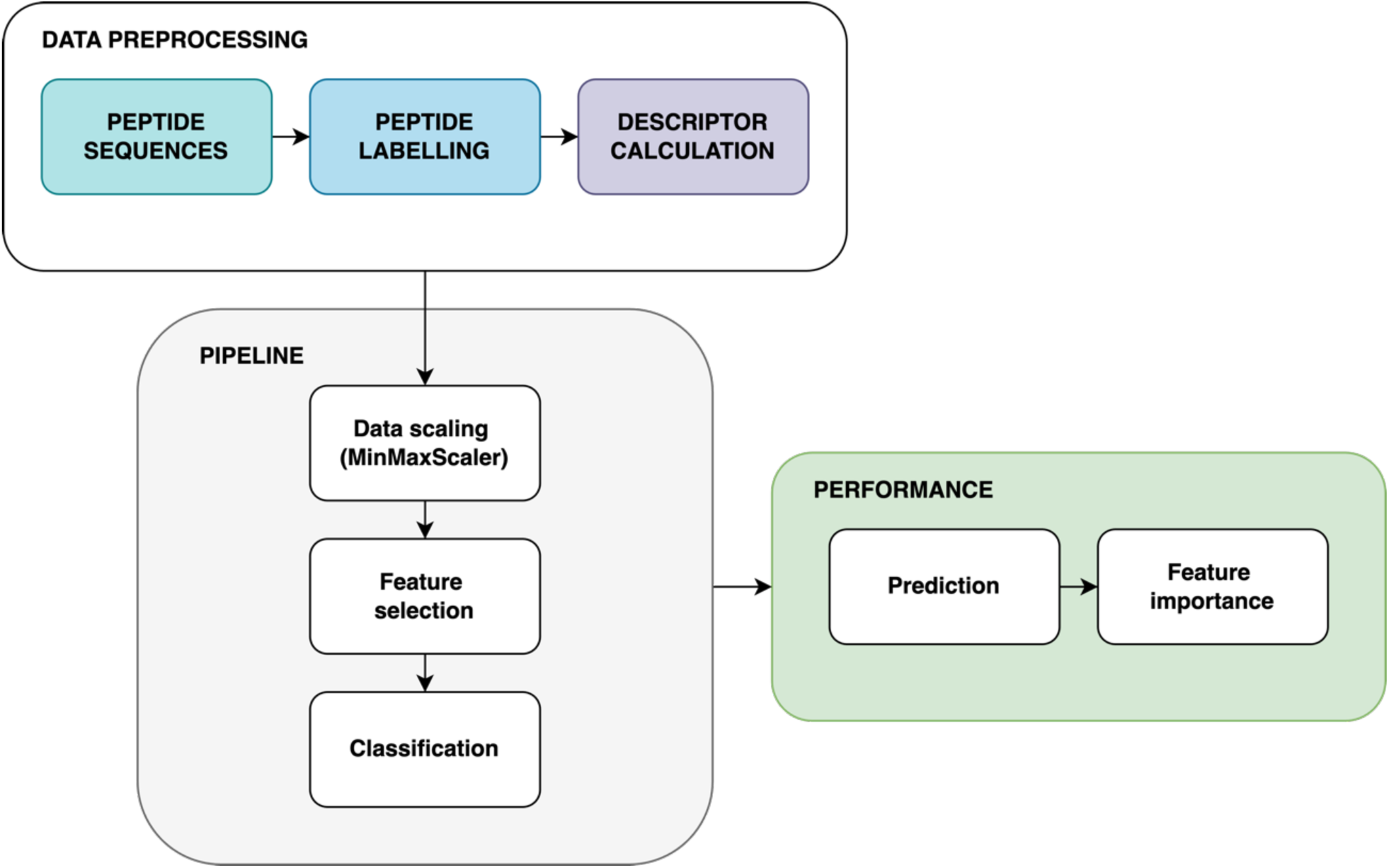
Data preparation and machine learning analysis scheme, showing peptide data pre-processing (**A**), the standardised classification pipeline (**B**) and performance evaluation and analysis (**C**).

The S-class metric ranged between 1 and -1, where 1 suggested perfect clustering, 0 indicated high overlap near the separation boundary, and -1 indicated poor clustering.

Results showed that all the S-clust scores were negative, confirming lack of separation. However, the t-SNE with perplexity set to 100 performed marginally superior (S-clust score = -0.06) whilst UMAP performed the worst (S-clust score = -0.11), as summarised in **Table 3**.

**Table 3.**
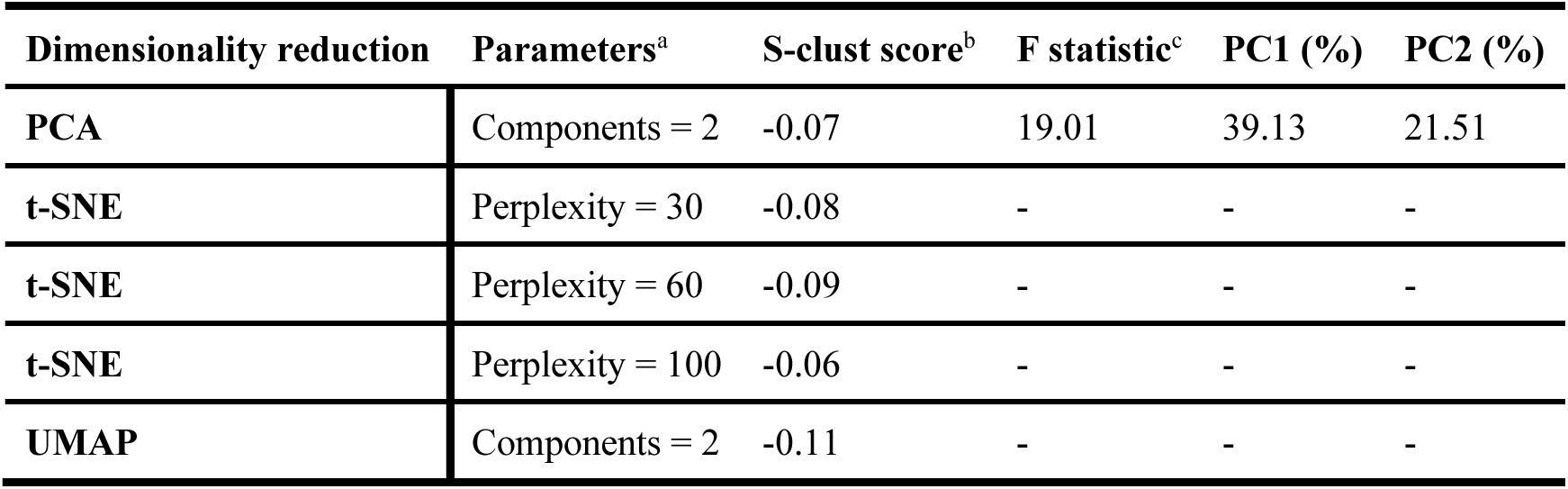
Summary of dimensionality reduction performance metrics, encompassing model parameters, Silhouette scores and F-statistic. challenging.

| Dimensionality reduction | Parameters<sup>a</sup> | S-clust score<sup>b</sup> | F statistic<sup>c</sup> | PC1 (%) | PC2 (%) |
| --- | --- | --- | --- | --- | --- |
| PCA | Components = 2 | -0.07 | 19.01 | 39.13 | 21.51 |
| t-SNE | Perplexity = 30 | -0.08 | - | - | - |
| t-SNE | Perplexity = 60 | -0.09 | - | - | - |
| t-SNE | Perplexity = 100 | -0.06 | - | - | - |
| UMAP | Components = 2 | -0.11 | - | - | - |

The lack of separation in descriptor dimensionality reduction suggested that the descriptors used in this study did not capture differences between peptide groups. This finding was consistent with the feature overlap already observed in peptide compositional and physicochemical property analyses. However, the statistical test on principal component 1 (PC1) in PCA analysis showed a high F-test score (F=19.01, **Table 3**), suggesting that underlying differences between the peptide classes exist but are not linearly separable. This observation indicated that more complex feature space boundaries may be present. Therefore, we have next used supervised decision tree classifiers to analyse our data.

### Classification of cell and brain penetrating peptides

Given a small dataset, imbalanced peptide class distribution and overlapping features we selected three tree-based classifiers: the classical Random Forest ^34^, Extremely randomised trees (ExtraTrees) ^35^ and extreme gradient boosting (XGBoost) ^36^. Tree-based classification algorithms were chosen based on their suitability to noisy data and ability to capture non-linear, more complex feature interactions. The Random Forest classifier is robust against overfitting of data ^34^, whilst XGBoost and Extra trees classifiers have been shown to be better suited for imbalanced and heterogenous datasets ^37^. A great advantage of decision tree-based algorithms is their simplicity and interpretability, which allowed us to extract feature importance values and their contribution to the model’s performance. To assess our model performance, we used standard metrics such as F1-score, confusion matrix, and precision-recall curves.

Our results showed comparable performance metrics of accuracy and F1-scores amongst classifiers, with the F1-scores of 0.68 for ExtraTrees and 0.66 for Random Forest and XGBoost, respectively (**Figure 7A**). The prediction of the best performing ExtraTrees classifier’s outcomes was summarised in confusion matrices (**Figure 7B-C**). The BPPs and CPPs showed 70% and 77% predicted recall values respectively, whilst non-BPPs showed only 12% recall. The low prediction of non-BPPs could be explained by our imbalanced dataset, where non-BPPs was a minority class.

**Figure 7.**
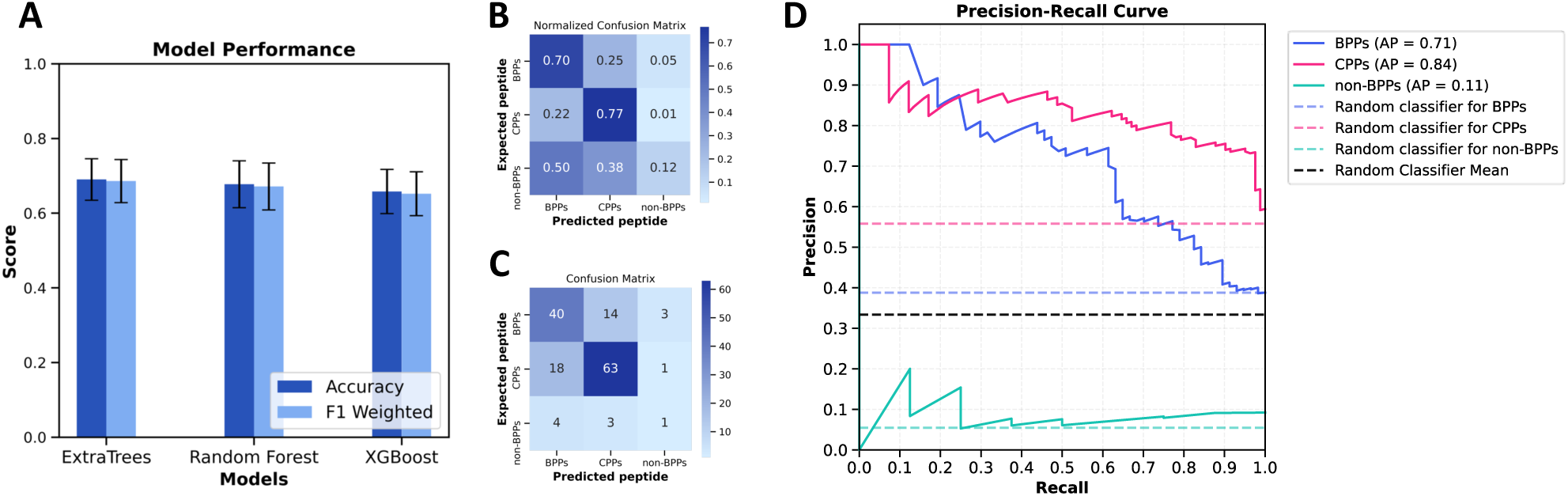
Multiclass classification of BPPs, CPPs and non-BPPs based on Modlamp peptide feature descriptors. **(A)** summarises the cross-validation average prediction score (F1-score) and accuracy metric of each classifier. Further performance analysis is summarised by confusion matrix in **(B)** and normalised confusion matrix **(C)**, whilst **(D)** indicates the precision-recall curve. AP = Average Precision.

The precision-recall analysis was consistent with the overall performance as indicated by F1-scores (**Figure 7D**, **Table 5)**. It showed that CPPs had the highest average precision score (AP=0.81), followed by BPPs (AP=0.71) whereas non-BPPs had the lowest AP score of 0.11. Both BPPs and CPPs recall were greater than random classifier, indicated by dotted lines as a threshold (Figure 7D). In contrast, non-BPPs showed nearly random precision-recall, suggesting that distinguishing non-BPPs from CPPs and BPPs was indeed

**Table 4.** Summary of classifier performance, where F1-score is an indicator of predictive performance of each classifier. The best performer and the model selected for further analysis is indicated by underline.

| Classifier | F1-score |
| --- | --- |
| <u>ExtraTrees</u> | <u>0.68</u> |
| Random Forest | 0.66 |
| XGBoost | 0.66 |

**Table 5.** Best classifier (ExtraTrees) classification performance scores.

|  | Precision | Recall | F1-score | Support |
| --- | --- | --- | --- | --- |
| **BPPs** | 0.65 | 0.70 | 0.67 | 57 |
| **CPPs** | 0.79 | 0.77 | 0.78 | 82 |
| **Non-BPPs** | 0.20 | 0.13 | 0.15 | 8 |
| **accuracy** | 0.71 | 0.71 | 0.71 | 0.71 |
| **macro avg.** | 0.54 | 0.53 | 0.53 | 147 |
| **weighted avg.** | 0.70 | 0.71 | 0.70 | 147 |

### Feature importance

Further analysis of the ExtraTrees classifier revealed which features contributed the most to differentiate CPPs, BPPs and non-BPPs. Our permutation feature importance (PFI) results identified pI, charge, and aromaticity as the strongest global predictors of peptide class separation (**Figure 8A**). A similar trend was observed in individual peptide class PFI scores (Figure 8B-C). However, CPPs had a hydropathy ratio as one of the top features whilst non-BPPs had an instability index. These results were consistent with previous observations, such as elevated cationic amino acid frequencies in CPPs and BPPs and greater aromaticity in CPPs (**Figure 1-2**). Interestingly, charge density was ranked as the least important feature in PFI in contrast to earlier observed sequence logo plots of amino acids (**Figure 2**). The molecular weight also appeared to be a less important feature, which can be explained by the similar lengths of peptides within CPPs and BPPs groups. Although non-BPPs were observed to be shorter peptides in our dataset, these results suggested that algorithm did not solely rely on peptide size differences but rather identified more meaningful physicochemical features such as pI and charge.

**Figure 8.**
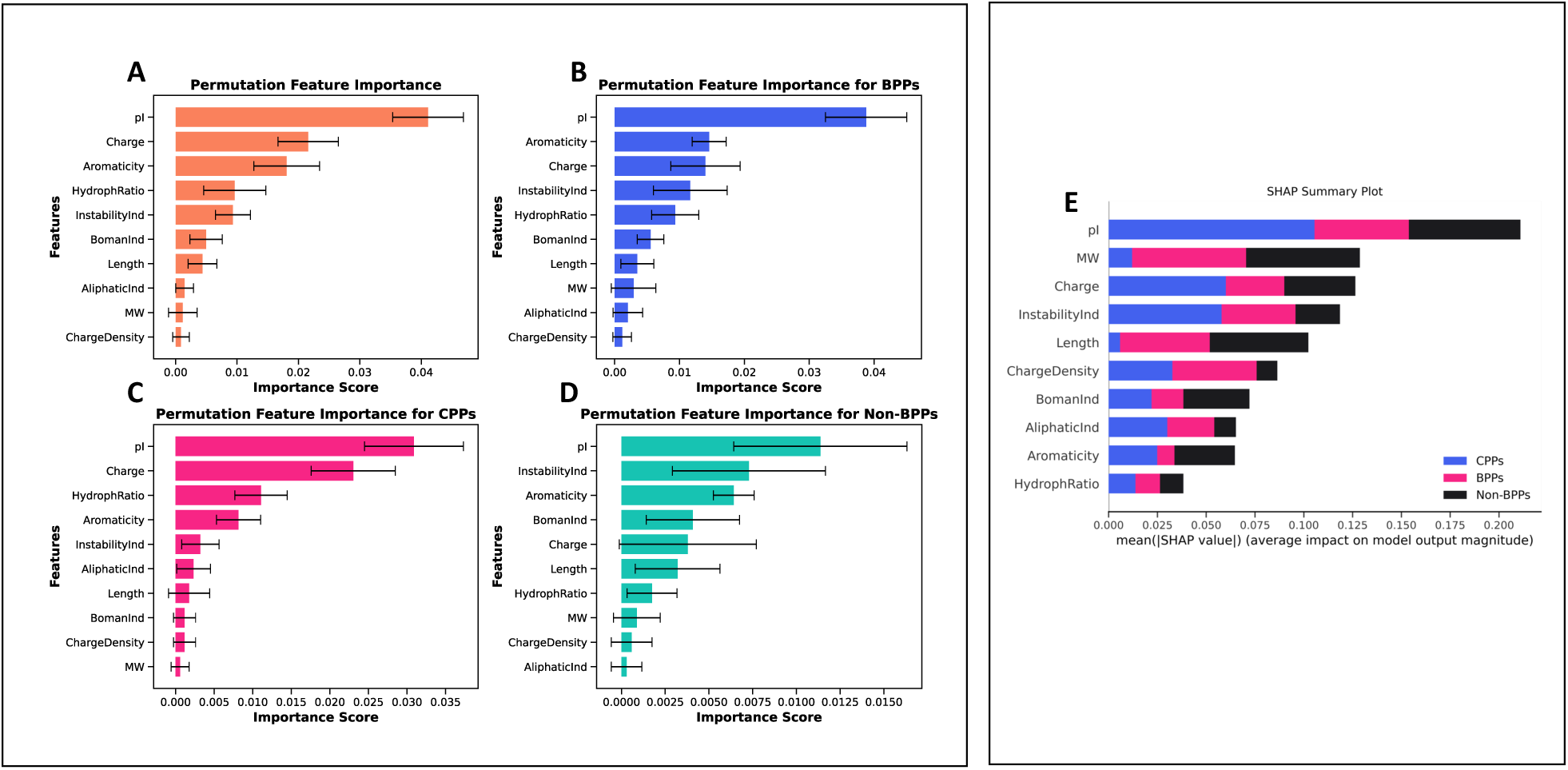
Feature importance (Modlamp descriptors) based on ExtraTrees classifier. (**A-D**) Permutation feature importance (PFI), where (**A**) is overall PFI and B,C,D are PFI’s of BPPs, CPPs and non-BPPs respectively. (**E**) SHAP feature importance.

However, it is key to note that PFI calculated the importance score by randomly shuffling the descriptors and then measured the classification performance penalties by correlating reduced model performance to greater feature importance. In contrast, the SHAP method calculated Shapley scores ^38^ based on the game theory framework, where all possible combinations of features are considered, and their global contribution to the model’s performance is estimated. Furthermore, permutation importance does not consider inter-feature correlation because of random shuffling, which could lead to misleading results, whilst the SHAP approach is not immune to the overfitting of the classifier as opposed to the permutation method, highlighting the limitations of both methods. The SHAP analysis results indicated, in agreement to PFI, that pI and charge were amongst the top important features, at least when comparing BPPs and CPPs (**Figure 8E**).

## Discussion

This study systematically explored the characteristic features of a literature-derived database of non-synthetic membrane-active peptides, categorised as BPPs, CPPs and non-BPPs. Here we hypothesised that cell-penetrating and brain-penetrating peptides despite their shared similarities such as interaction with cellular membranes through internalisation and translocation mechanisms, have distinct physicochemical differences. Indeed, our results demonstrated that, in agreement with the literature, these peptides are predominantly cationic and amphipathic ^15,39^. However, we identified subtle but important differences in amino acid composition between BPPs and CPPs. The CPPs had predominantly cationic arginine residues, whilst BPPs consisted of mixed arginine/lysine residues and more distributed cationic charge densities compared to CPPs. Amino acid positional analysis (**Figure 4A-B**) further revealed that CPPs had cationic charge clustering at their core/C-terminus whilst BPPs had more evenly spread cationic charge. This is in agreement with a molecular dynamics simulation study showing that distributed cationic charge on peptide sequence is more favourable for permeating cellular membranes due to reduced membrane distortion compared with clustered cationic charge, as demonstrated by Povilaitis et al ^40^. Notably, non-BPPs had relatively neutral and hydrophobic amino acids and were smaller than CPPs and BPPs which accords with literature reports linking higher hydrophobic content within peptides to greater cytotoxicity ^21^.

Brain-penetrating peptides showed lower isoelectric point than CPPs which could be explained by the enrichment of arginine (pKa ≈ 12 ^41^) in CPPs whilst BPPs had a mixture of lysine (pKa ≈ 10 ^41^) and arginine. Since pI values close to physiological pH render peptides less cationic ^29^, this characteristic may contribute to peptide’s biocompatibility whilst highly cationic peptides often lead to cell lysis and are better suited as antimicrobial agents ^42,43^. BPPs also exhibited lower instability index, which measures dipeptide composition ^29^, suggesting that these peptides were more stable in an aqueous environment. Interestingly, we observed that BPPs at their core were more hydrophobic than CPPs, suggesting that the location cationic and hydrophobic amino acids along the peptide sequence may be important when designing brain-penetrating peptides or peptide-mimicking agents.

Whilst composition and physicochemical properties revealed differences between peptide groups, dimensionality reduction did not separate these classes into distinct clusters, suggesting a more complex structural diversity of peptides and more nuanced physicochemical differences. Non-linear dimensionality reduction algorithms improved peptide class separation only modestly as indicated by higher S-clust scores, suggesting that physicochemical correlation was interlinked, perhaps in a non-linear fashion.

This observation can be explained by our amino acid composition analysis, where most amino acids were similar amongst BPPs, CPPs, and non-BPPs. Alternatively, the choice of physicochemical descriptors in this study may not fully captured key aspects of peptide differences as descriptor performance depends on how well they represent peptide sequence-function relationships ^44^. To better understand the differences between BPPs, CPPs and non-BPPS, we employed supervised decision tree machine learning algorithms, where each peptide group was labelled. The results indicated that all tested tree-based classifiers, especially the ExtraTrees algorithm, could distinguish CPPs from BPPs but not non-BPPs. The latter was likely due to the underrepresentation of non-BPPs within the dataset. Indeed, imbalanced datasets are a common obstacle in machine learning ^45^, and new methods are continuously being investigated to alleviate this ^46^. Brain-penetrating peptides are particularly underrepresented in literature as shown by our dataset and a recent meta-analysis of brain penetrating peptides ^15^. However, despite this limitation we extracted important features that contributed to the classification of peptides. Generally, these features, such as pI, charge and aromaticity, aligned with our data analysis results, suggesting that the model identified and used relevant descriptors to distinguish between different peptides classes. The latter result may be important to consider when designing brain-penetrating peptides or even their synthetic alternatives.

It is important to discuss limitation of our classification study. Firstly, our classification model was based on a small dataset of CPPs, BPPs and non-BPPs. This could explain the limited classification power in the non-BPP class. Furthermore, this study deliberately used a small amount of easily interpretable descriptors. Indeed, the choice of descriptors can significantly influence the performance of machine learning classifiers. Feature engineering is often employed to select or combine multiple complex descriptors representing peptides ^47^, often leading to high dimensional descriptor spaces, which may improve classification performance but are much harder to interpret. Our rationale behind the descriptors in our study was that they could be easily interpretable and translatable into novel peptide or polymer synthesis. Alternative types of descriptors could be considered for future work. One example would be converting the peptides to their corresponding molecular representations, as described in the molecular “cliques” approach by Bernard et al. ^48,49^. A more recent approach addressed this with a genetic algorithm approach to automatically identify most optimal descriptor subsets for antimicrobial peptides which minimised bias of predefined descriptor sets ^44^. Future studies should also consider peptides’ secondary structure. For example,

Liu et al. showed that molecular dynamic simulation could be used to extract antimicrobial peptide descriptors linked to their 3-dimensional (3D) shape ^50^. Such an approach is time-consuming and limited by the accuracy of molecular dynamic simulations. However, it is important to consider the conformational behaviour of peptides. For the peptide to form a secondary structure, typically, it requires a driving environment, such as hydrophobicity. As an example, the latter was demonstrated by Gong *et al.*, ^51^ where some CPPs were shown to remain in a random coil structure when interacting with the cell membrane. However, the secondary structure was more prominent when inserted within a hydrophobic phospholipid. Hence, binding and insertion could be linked to different peptide shape characteristics. The latter suggests that looking beyond simple physicochemical descriptors could be considered in the future, especially when designing novel compounds.

Finally, different machine learning approaches, such as neural networks, could be used. Pandey *et al.* ^52^ used a Kernel Extreme Learning Machine (KELM), a type of neural network, to classify CPPs with over 80% accuracy, using compositional descriptors. However, the limiting factor, specifically in neural networks, is the quality and quantity of data. For example, cell-penetrating and antimicrobial peptide databases are more widely available, with larger datasets than BPPs. A more advanced BPP predictor was described by Ma *et al.* ^53^where a transformer-based convolutional neural network was used. There, too, the authors only had a limited number of peptides, which were augmented by artificial data augmentation methods. Such a model shows promising results i.e., 98% accuracy in prediction, which showcases the potential of machine learning applications within the field. There are, however, some caveats: results are harder to interpret and suffer from the necessity of data augmentation, which makes it overall and much harder to extract meaningful feature importances.

In summary, machine learning is showing great potential in shaping the understanding of membrane-active peptide characteristics. So far, however, the remains to be limited by good quality and experimentally validated datasets. The current literature describes many different peptides grouped into various classes, such as antimicrobial, cell-penetrating, membrane-active, brain-penetrating and shuttle peptides ^54^. Naturally, there is some overlap between these groups, which might perhaps make these peptides not easily or not at all distinguishable, as seen in our study. Generally, most membrane-active peptides exhibit cationic and amphipathic properties with overlapping physicochemical space ^55^. Furthermore, a much smaller number of peptides has been experimentally tested for blood-brain barrier penetration ^15^, and often, in-vitro tests show variability in methodology and experimental settings, as seen in common brain-penetrating peptide open-source databases ^56^. Even given these caveats, we showed that it is possible and useful to extract important features of different classes of peptides. In the future, perhaps it would be more promising to employ experimental data augmentation of peptides, such as BPPs, by utilising high-throughput in-vitro assays or even a completely different approach, for example, by using efficient and inexpensive synthetic polymer chemistry as a platform to indirectly assess various properties of the peptides.

## Conclusions

This study investigated the features of CPPs, BPPs and non-BPPs using machine learning multiclass classification. Despite limited datasets and descriptor space, we identified characteristic features of these groups of peptides. BPPs had a more balanced cationic charge and charge density along the peptide sequences when compared to CPPs, which were predominantly cationic. In contrast, non-BPPs were markedly less cationic and more hydrophobic. Random forest, ExtraTrees and XGBoost decision tree classifiers were able to classify CPPs and BPPs to some extent. non-BPPs, however, proved to be harder to classify using our machine-learning approach. Nevertheless, we extracted some important features contributing to peptide classification. The latter allowed us to refine our focus on the most important features when distinguishing BPPs, CPPs and non-BPPs. This demonstrates that machine-learning classification can be used as a tool to extract meaningful physicochemical properties of the peptides from limited and imbalanced datasets. In the future, more comprehensive descriptors, larger datasets backed up by in-vivo data and more advanced machine-learning algorithms should be explored.

Finally, this study allowed us to understand better the complex nature of short-sequence peptides that penetrate and permeate the cells. The machine-learning research and application fields are expanding rapidly. However, significant limitations, such as small and inconsistent datasets, must first be overcome to fully benefit from these approaches. Future work could include high throughput in-vitro blood-brain barrier assays to augment the limited datasets and address their imbalances. Overall, this study attempted to work towards understanding the physicochemical properties of short brain penetrating peptides. These findings can now be used as a reference guide to instruct the rational design of synthetic peptide-mimicking polymers and assess their biological relevance.

## Acknowledgements

VK was funded by the Medical Research Council Interdisciplinary Biomedical Research Doctoral Training Partnership (MR/N014294/1).

## Supplementary Data

Peptide database of brain-penetrating peptides (BPP), cell-penetrating peptides (CPP) and nonbrain-penetrating peptides (Non-BPP)

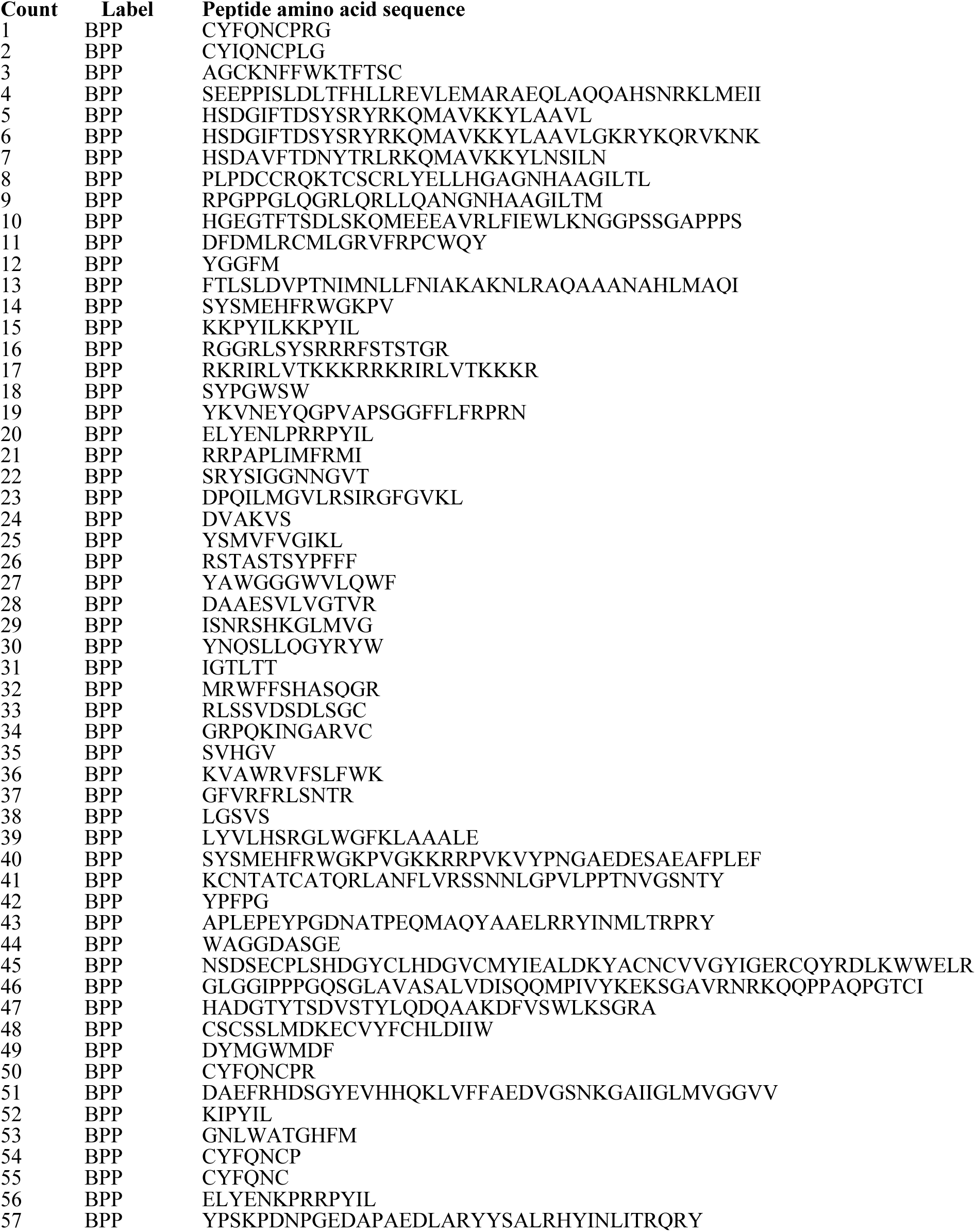
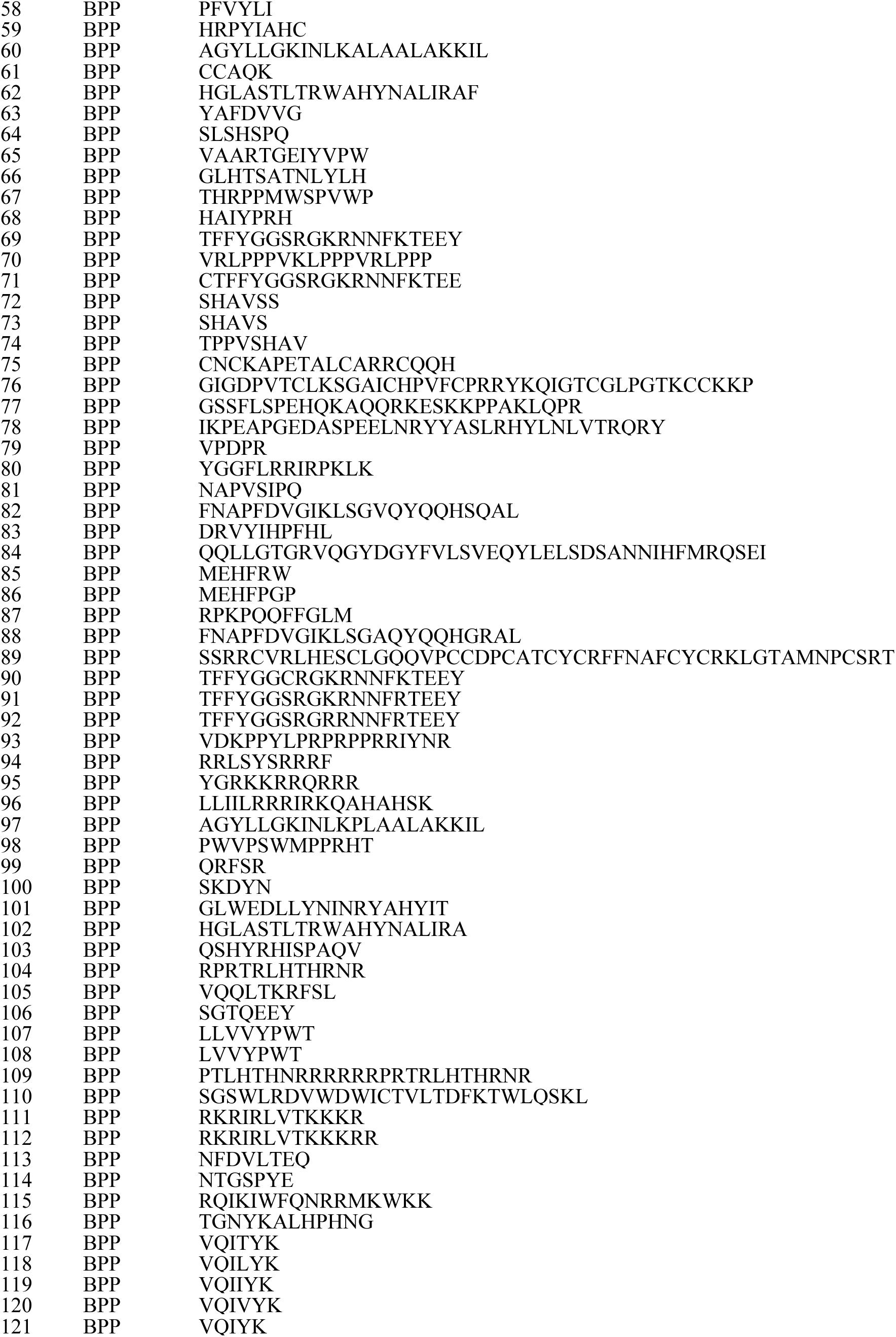
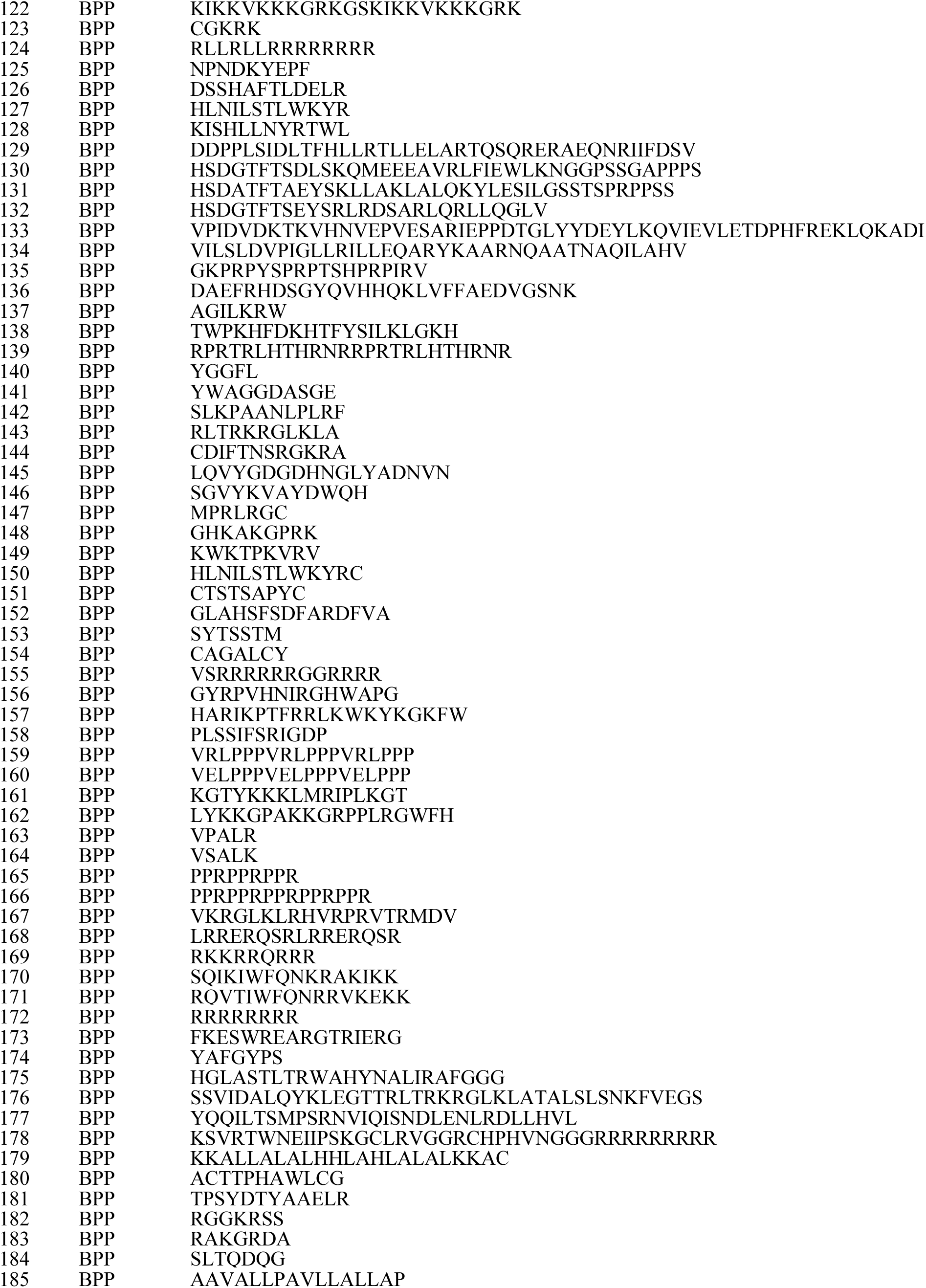
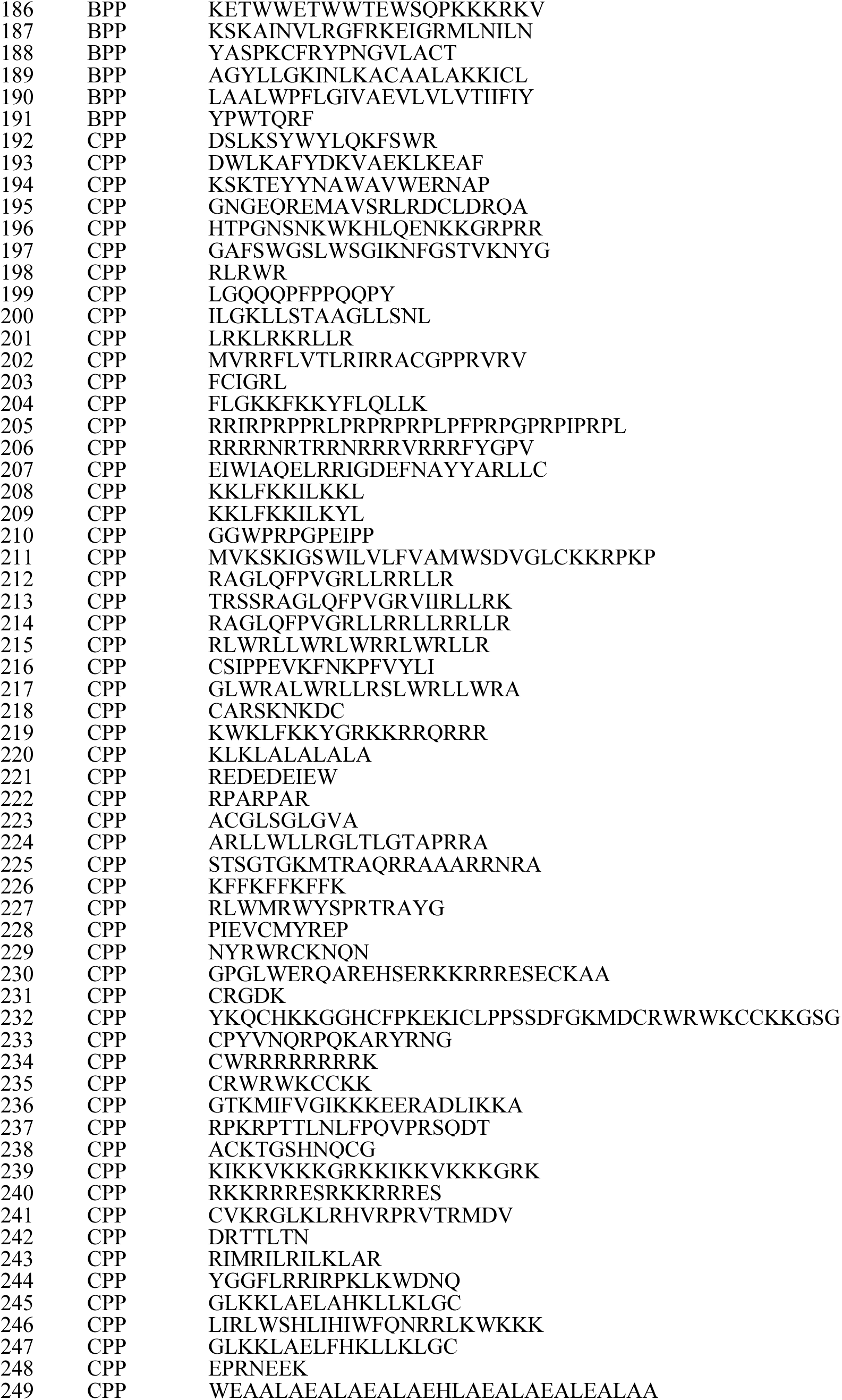
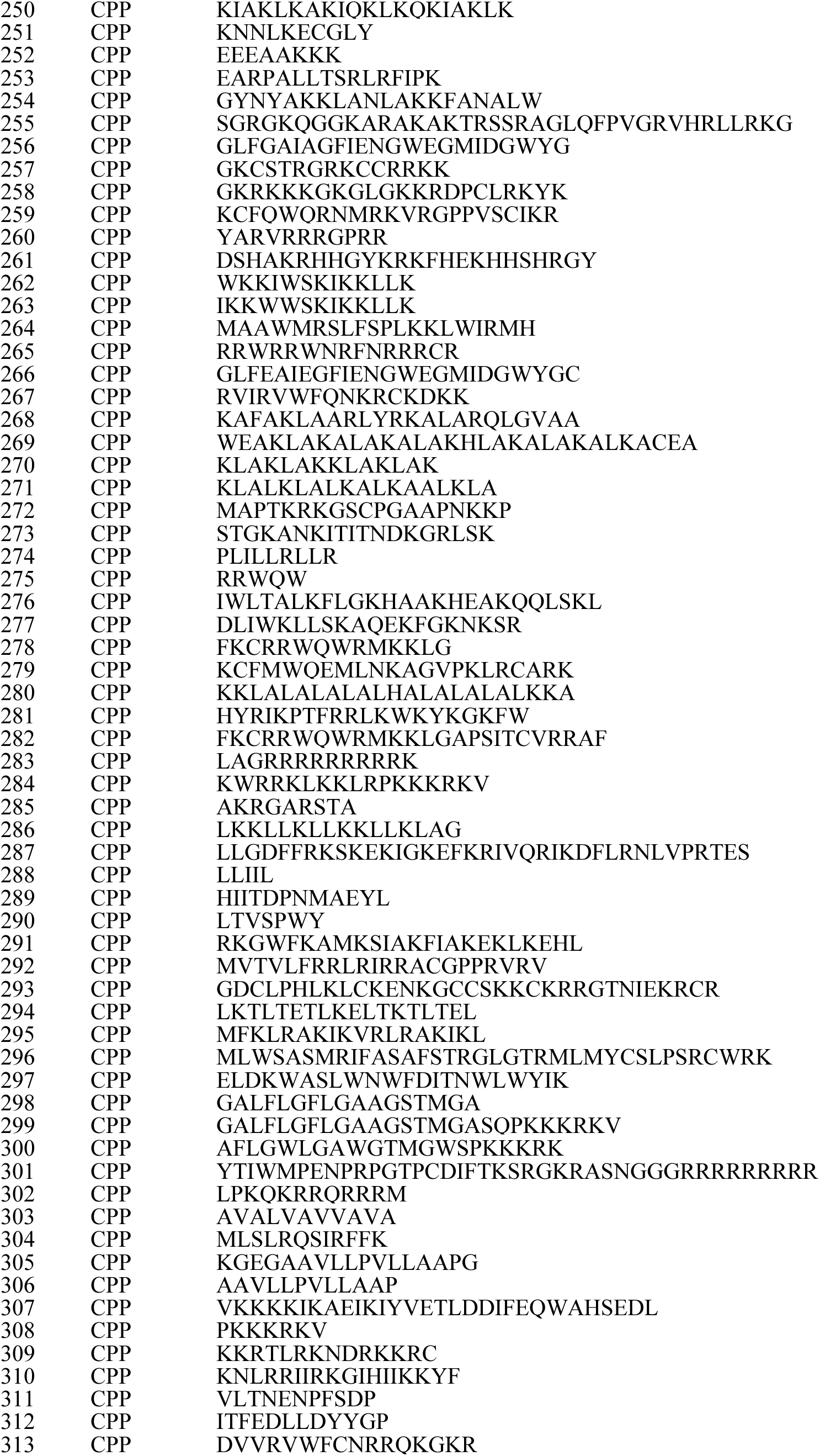
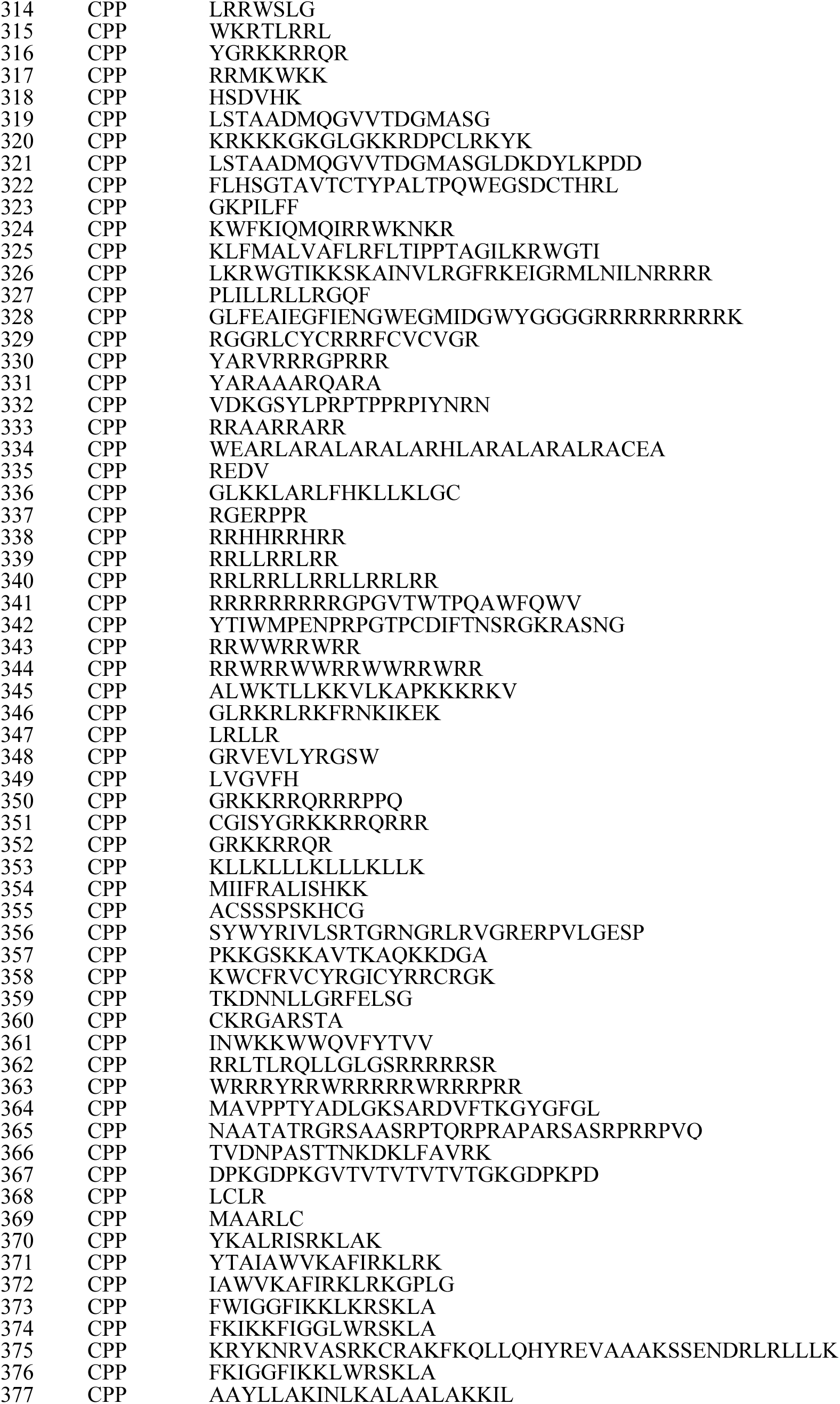
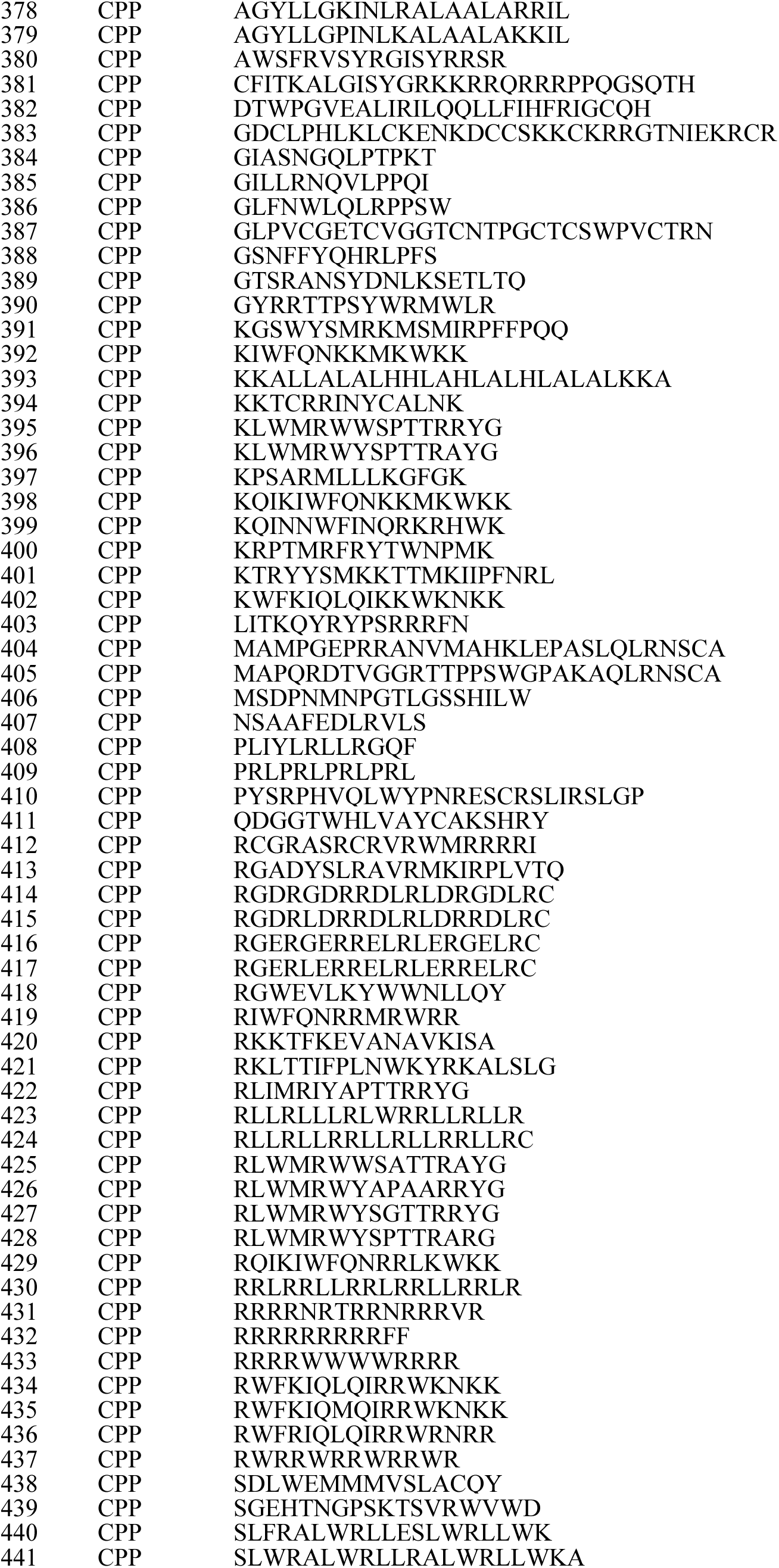
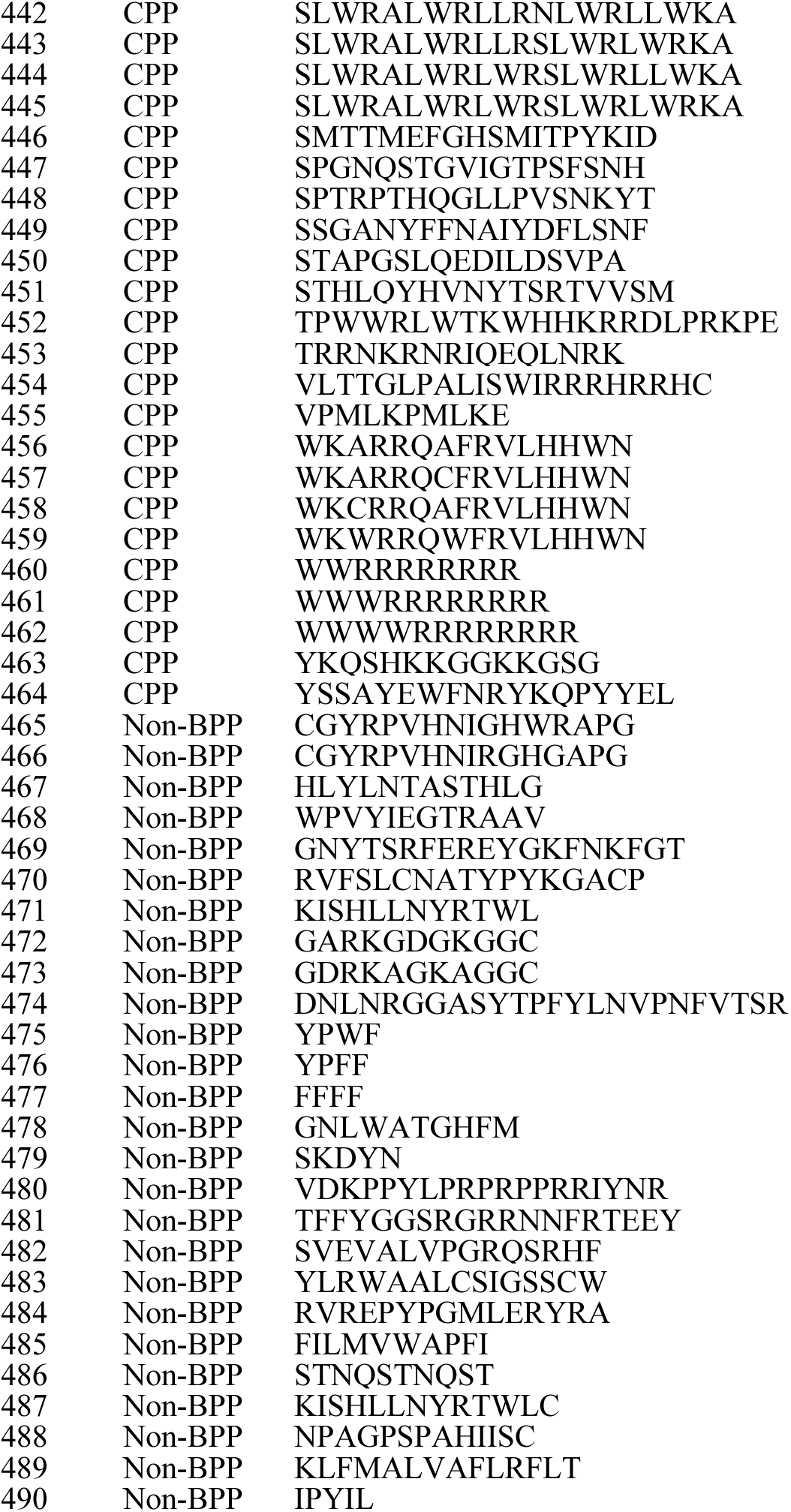
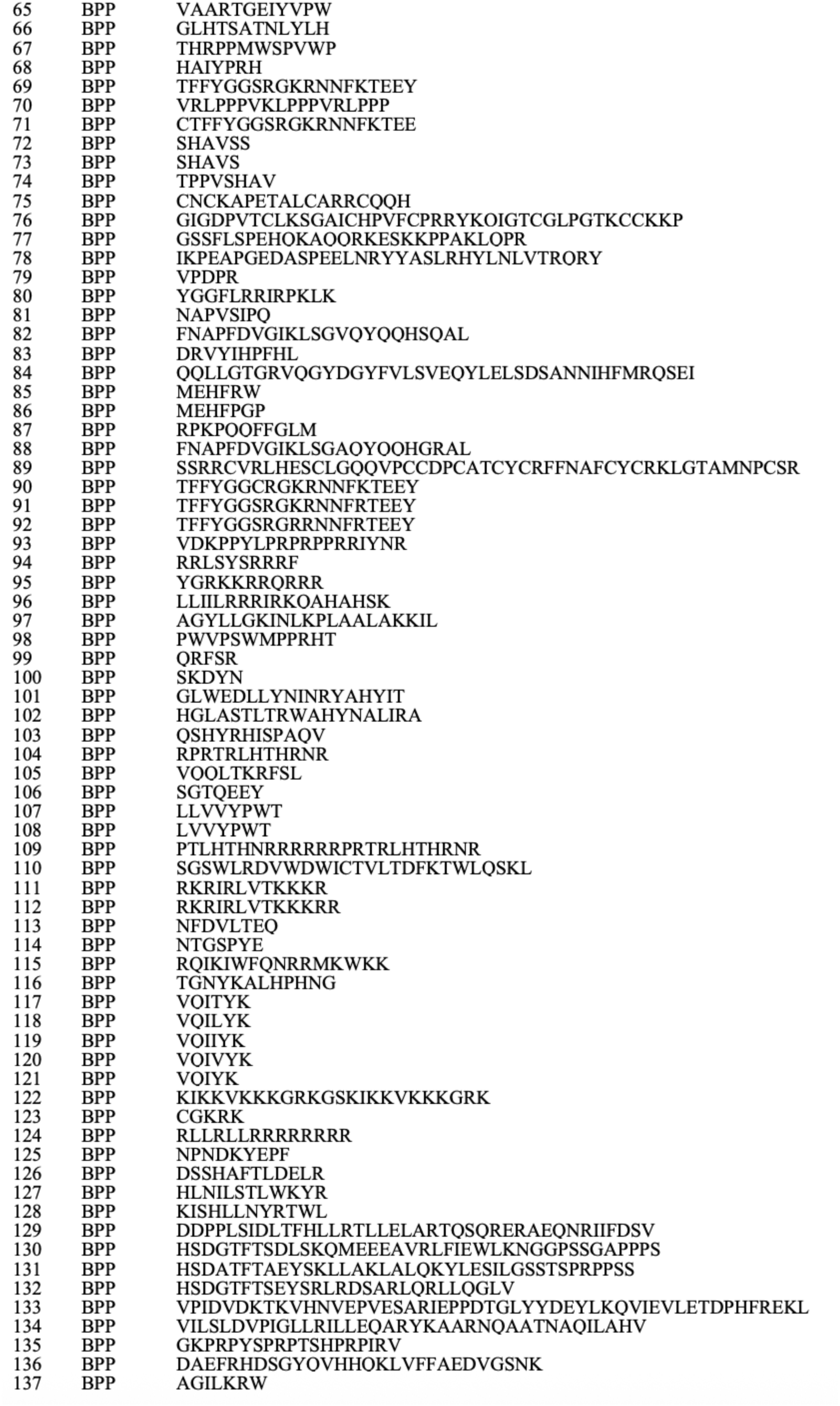
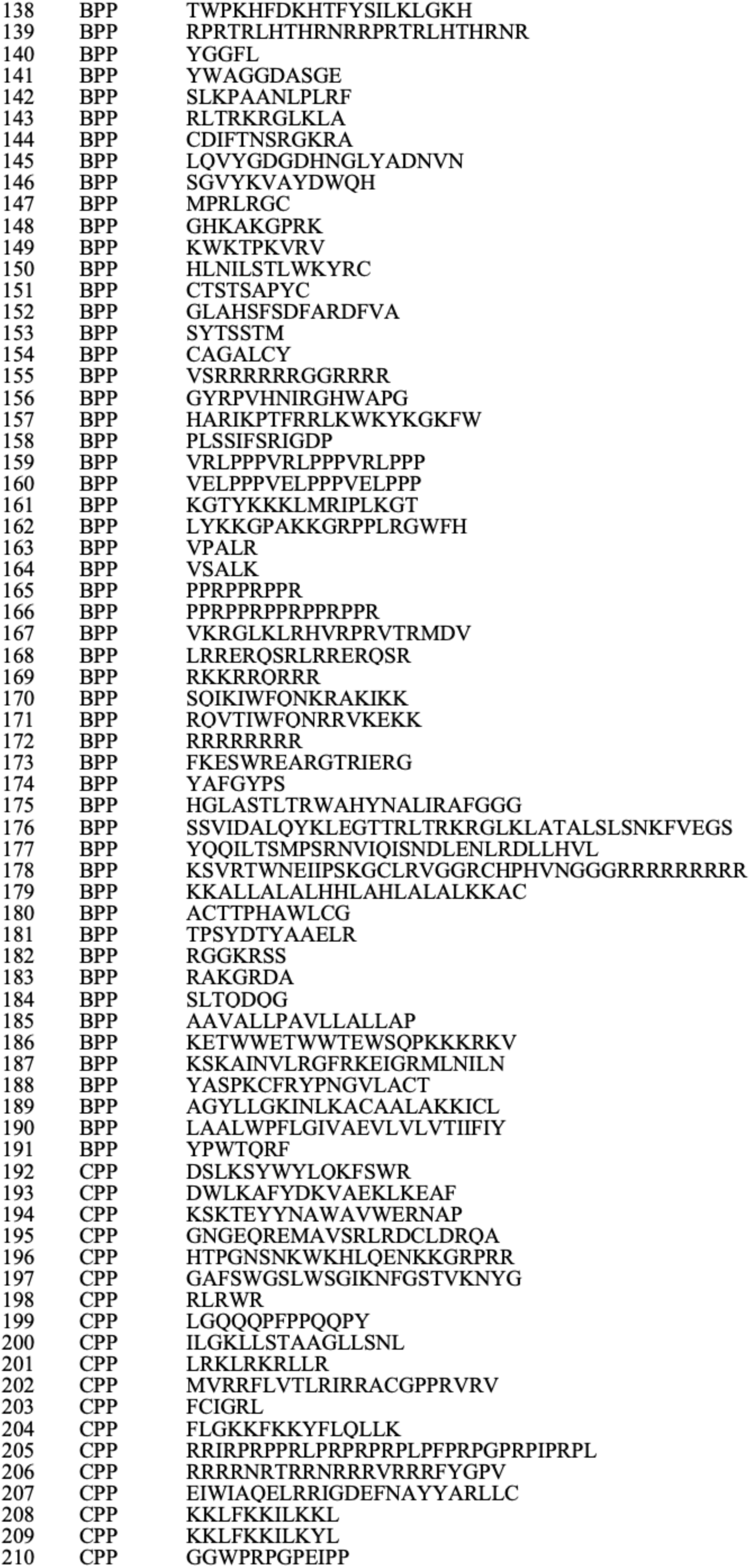
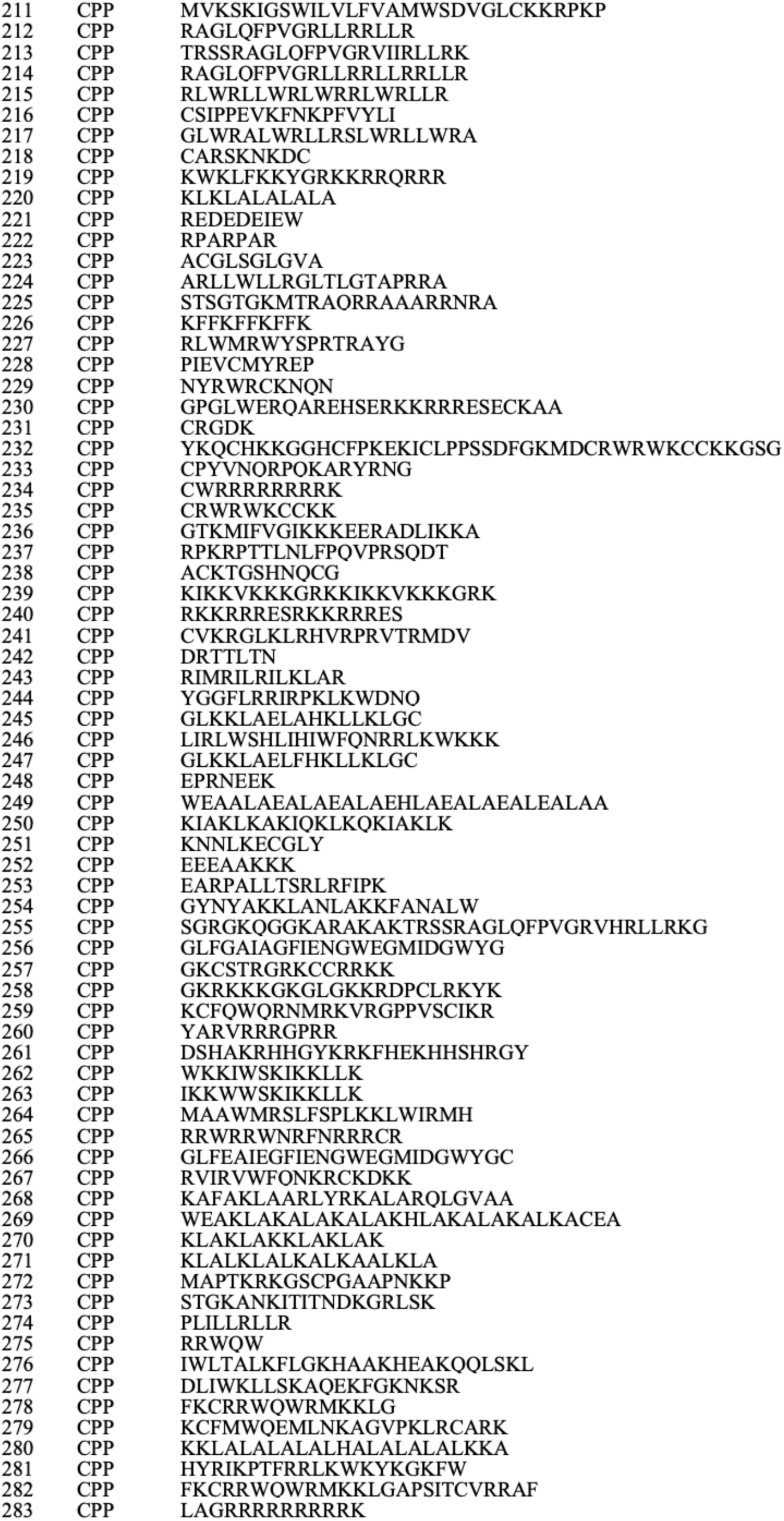
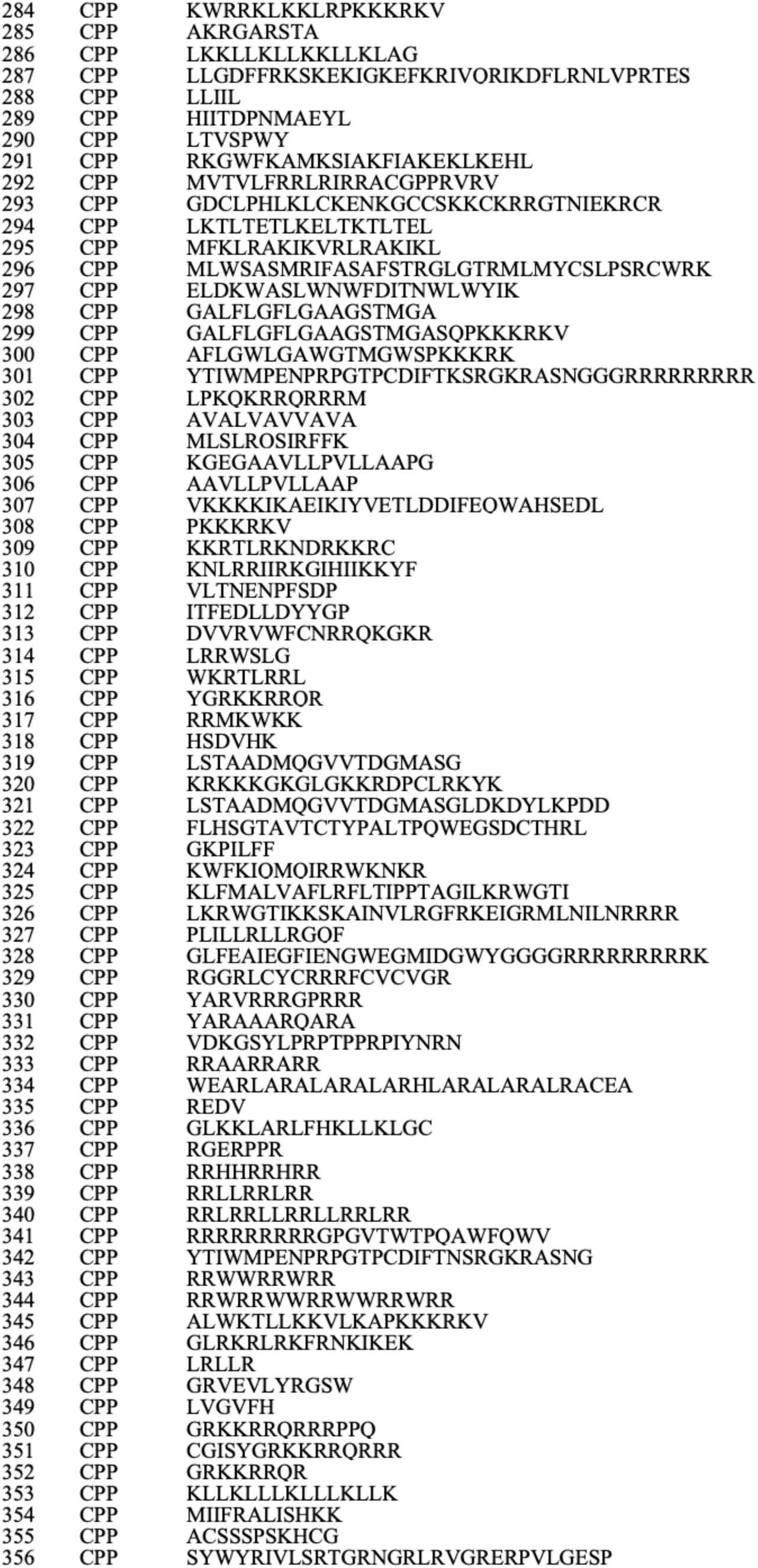
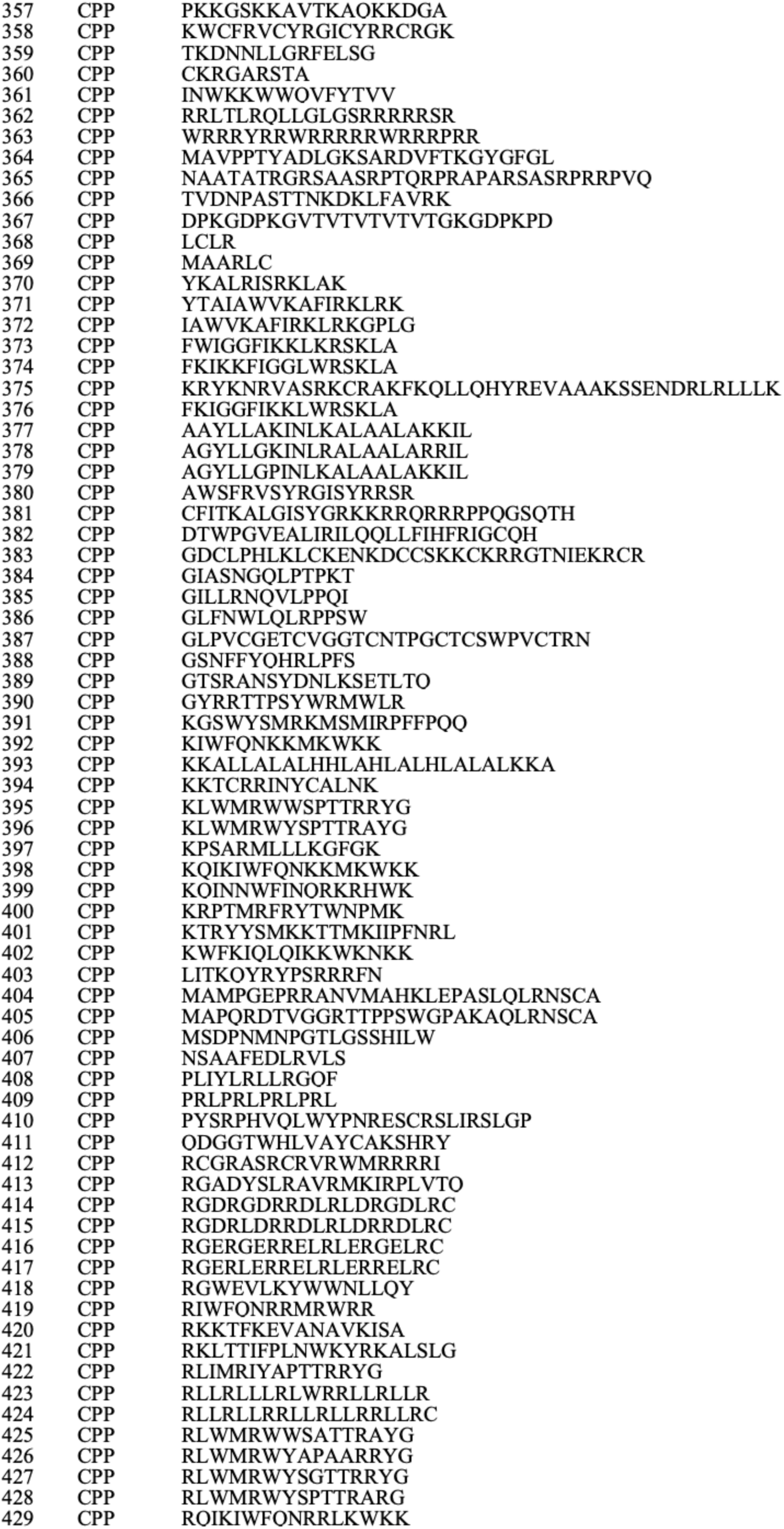
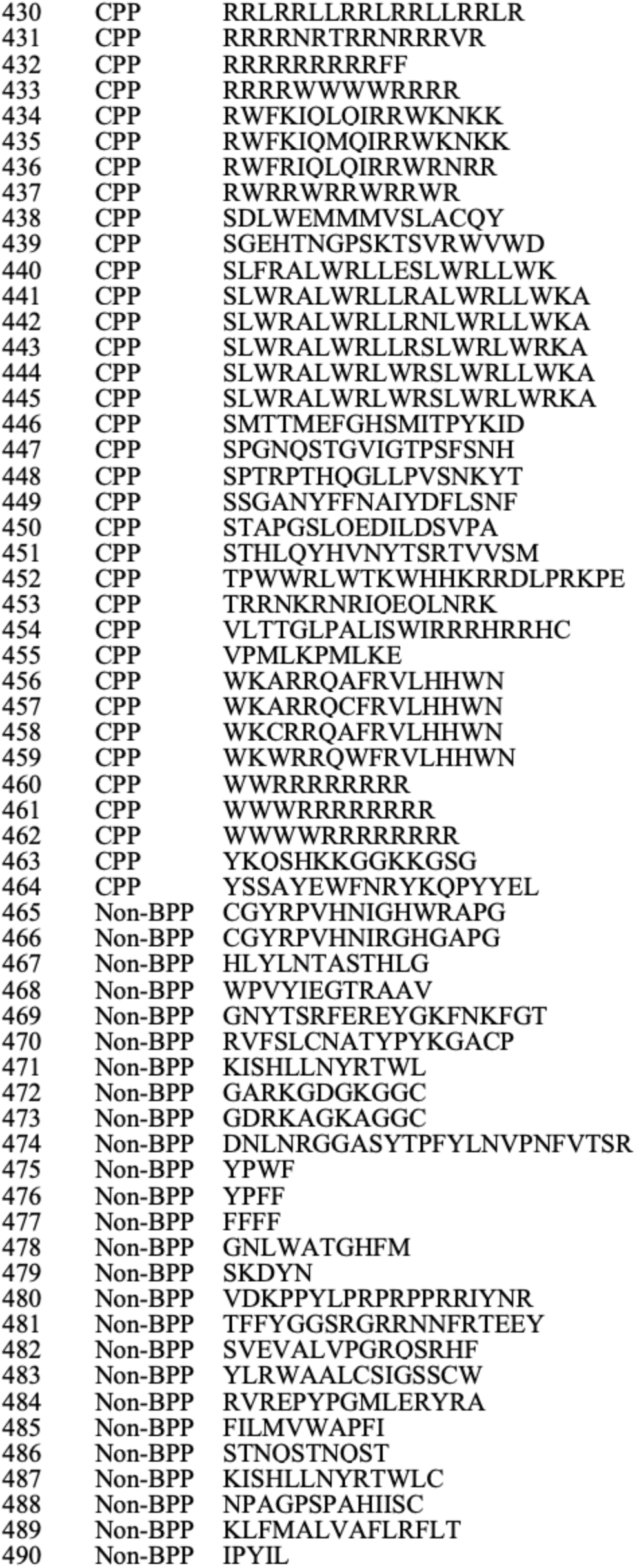

